# Emergence of isochorismate-based salicylic acid biosynthesis within Brassicales

**DOI:** 10.1101/2025.03.03.641121

**Authors:** Kunqi Hong, Ying Tang, Linda Jeanguenin, Wenshang Kang, Yongliang Wang, Lu Zuo, Pengyue Li, Jingjng He, Wanqing Jiang, Ruidong Huang, Hidenori Matsui, Yiming Wang, Hirofumi Nakagami, Bo Li, Xia Li, Kabin Xie, Kenji Fukushima, Liang Guo, Xiaowei Han, Fumiaki Katagiri, Motoyuki Hattori, Kenichi Tsuda

**Affiliations:** National Key Laboratory of Agricultural Microbiology, Hubei Hongshan Laboratory, Hubei Key Laboratory of Plant Pathology, College of Plant Science and Technology, Huazhong Agricultural University, Wuhan, China; Department of Plant and Microbial Biology, University of Minnesota – Twin Cities, St Paul, MN 55108, USA; National Key Laboratory of Crop Genetic Improvement, Hubei Hongshan Laboratory, College of Plant Science and Technology, Huazhong Agricultural University, Wuhan, China; Plant Proteomics Research Unit, RIKEN Center for Sustainable Resource Science, Yokohama 230-0045, Kanagawa, Japan; Graduate School of Environmental and Life Sciences, Okayama University, Okayama 700-8530, Japan; State Key Laboratory of Agricultural and Forestry Biosecurity, College of Plant Protection, Nanjing Agricultural University, Nanjing,210095, China; Basic Immune System of Plants, Max-Planck Institute for Plant Breeding Research, Cologne, 50829, Germany; Center for Frontier Research, National Institute of Genetics, 1111 Yata, Mishima, Shizuoka 411-8540, Japan; State Key Laboratory of Genetic Engineering, Collaborative Innovation Center of Genetics and Development, Department of Physiology and Neurobiology, School of Life Sciences, Fudan University, Shanghai 200438, China

**Keywords:** Brassicaceae, Brassicales, evolution, plant defense phytohormones, salicylic acid

## Abstract

Salicylic acid (SA) is a major defense phytohormone. In *Arabidopsis thaliana*, the isochorismate (IC) pathway is the primary route for pathogen-induced SA biosynthesis. First, the IC synthase (ICS) catalyzes the isomerization of chorismate to IC in chloroplasts. Second, the chloroplast-localized MATE transporter EDS5 appears to transport IC from chloroplasts to the cytosol. Cytosolic IC is then further converted to SA via the GH3 amino acid-conjugating enzyme PBS3. While this pathway is genetically well-characterized in *A. thaliana*, its evolutionary origin and conservation remain controversial. In this study, through comprehensive phylogenetic, structural, and functional analyses, we demonstrate that the IC pathway emerged within the Brassicales order in a time span between the divergence of *Carica papaya* and *Capparis spinosa*. The evolution of the IC pathway was driven by three key adaptations during the time span: (1) enhancement of ICS activity, (2) neofunctionalization of *EDS5* after duplication of its ancestral gene, and (3) evolution of a *PBS3*, whose activity is specialized for glutamate-conjugation to IC. Structural modeling and functional assays reveal that an enhanced salt bridge network in ICS enhanced its activity. One of the duplicated genes, EDS5, acquired key amino acid substitutions in the C-lobe, which likely contributed to the *EDS5* neofunctionalization. In addition, the functional *PBS3* clade, including *A. thaliana PBS3*, is restricted to a Brassicales clade. Taken together, this study addresses the evolutionary trajectory of IC-based SA biosynthesis.

## INTRODUCTION

Salicylic acid (SA) is a phytohormone that regulates various plant physiological processes, including immunity, abiotic stress tolerance, and growth (1–3). Among these roles, SA is particularly prominent in plant immunity. Mutant plants deficient in SA biosynthesis or signaling exhibit compromised disease resistance (4, 5). Additionally, exogenous application of SA or its synthetic analog enhances plant resistance against pathogens, a strategy that has been utilized in agricultural practices to improve crop performance (6–8).

In angiosperms, SA biosynthesis occurs via two distinct pathways that share the common precursor chorismate: the phenylalanine ammonia-lyase (PAL) and the isochorismate (IC) pathways. In the PAL pathway, phenylalanine, presumably produced from chorismate, is converted to *trans*-cinnamate by PALs (9). Cinnamate is subsequently converted into benzoic acid by Abnormal Inflorescence Meristem 1 (AIM1) and then into SA by yet-identified benzoic acid-2-hydroxylases (BA2Hs) (10). The PAL pathway is supported by fragmentary evidence from studies using different plant species. For instance, *Arabidopsis thaliana pal* quadruple mutant plants exhibited a 50% reduction in pathogen-induced SA accumulation (11), and rice *pal6* mutants showed a 60% decrease in basal SA levels in roots (12). Knockdown of *ZmPAL* expression resulted in decreased SA accumulation in maize (13). *AIM1* was required for basal SA accumulation in rice leaves (14). In addition, isotope tracing experiments have supported the existence of the PAL pathway in *Nicotiana tabacum* (15). However, a recent study showed that feeding *A. thaliana* with ^13^C_6_-Phe resulted in the incorporation of the ^13^C label, not into SA but instead its isomer 4-hydroxybenzoic acid (16). Although BA2H activity has been detected in *N. tabacum* (15), no genes responsible for BA2H have been identified in any plant species. Thus, our knowledge of the PAL pathway in plants is incomplete.

In contrast, the IC pathway has been genetically elucidated in *A. thaliana*. In this pathway, the IC synthase (ICS) catalyzes the isomerization of chorismate to IC within chloroplasts (17). The Glycoside Hydrolase 3 (GH3) family protein AvrPphB Susceptible 3 (PBS3) catalyzes the conjugation of glutamate to IC in the cytosol, forming IC-9-glutamate, which is subsequently converted to SA either spontaneously or by ENHANCED PSEUDOMONAS SUSCEPTIBILITY 1 (EPS1) (18, 19). The MATE (multidrug and toxin extrusion) transporter ENHANCED DISEASE SUSCEPTIBILITY 5 (EDS5) localizes to the chloroplast membrane (4, 20). Although EDS5 is hypothesized to transport IC from chloroplasts to the cytosol, direct evidence is still lacking. However, the observation that chloroplast-targeted PBS3 restored SA biosynthesis in *eds5* mutant plants supports this hypothesis (19). Collectively, ICS, EDS5, and PBS3 constitute essential components of the IC pathway in *A. thaliana*, with EPS1 serving as an additional modulator. The IC pathway is considered the primary route for SA biosynthesis in *A. thaliana*. Once synthesized, SA is metabolized into SA 2-*O*-β-D-Glucoside (SAG) and salicylate glucose ester, which typically accumulate at five- to tenfold higher levels than SA itself in plants (21, 22).

The conservation of the IC pathway across plant species remains controversial. Silencing of *ICS* genes in soybean and cotton plants resulted in decreased SA levels (23, 24), and both the PAL pathway and *ICS* gene likely contribute to SA biosynthesis in maize, as evidenced by the incorporation of labeled phenylalanine into SA glucoside and the reduction of PAL activity in *ics1* mutant plants (25). However, genetic mutations in *ICS* genes did not affect SA levels in rice or barley (26, 27). A recent study proposed that the IC pathway evolved in the most recent common ancestor (MRCA) of land plants, given the presence of homologs of *ICS*, *EDS5*, and *PBS3* in land plant genomes (28). The *A. thaliana* genome encodes two *ICS* genes (*ICS1* and *ICS2*), with *ICS1* being primarily responsible for pathogen-induced SA accumulation, though *ICS2* also contributes (17, 29). Additionally, IC serves as a precursor of phylloquinone, an essential electron acceptor in photosynthesis (30). Thus, *ICS* genes are likely crucial for maintaining proper photosynthesis. Indeed, *A. thaliana ics1ics2* double and *vte6* mutant plants deficient in phylloquinone production showed severely compromised growth (29, 31). These observations suggest that the evolutionary conservation of *ICS* genes does not necessarily indicate their involvement in the IC pathway for SA biosynthesis.

*A. thaliana* possesses a homolog of *EDS5* (*EDS5H*), which is not required for SA biosynthesis (32). The biological functions of EDS5H are unknown, and EDS5 function in plant species beyond *A. thaliana* remains unexplored. Additionally, although homologs of *PBS3* have been identified across various angiosperm genomes (33), *PBS3* (*GH3.12*) belongs to a large *GH3* gene family, necessitating systematic evolutionary and functional analyses to determine its conservation. *EPS1*, on the other hand, appears to be conserved exclusively in Brassicaceae (18). Collectively, the evolutionary conservation of the IC pathway across plants remains unresolved.

Because SA plays important roles in responses to biotic and abiotic factors, understanding the evolution of the IC pathway is critical for elucidating how plants have adapted to their environment over evolutionary timescales. Insights into the IC pathway evolution could also facilitate the development of crop varieties with enhanced resilience to environmental stresses and pathogens. Ohno’s hypothesis (34) suggests that gene duplications drive the emergence of novel traits, fostering complexity and species diversity (35, 36). Whole genome duplication (WGD) is a key mechanism in plant speciation and evolution (37, 38). Genomic analyses have identified multiple WGD events in plants. For instance, the At-β WGD occurred within Brassicales after the divergence of *Carica papaya* (39). The evolution of SA biosynthetic enzymes likely involved gene duplication, neofunctionalization, and gene co-option. However, significant knowledge gaps remain regarding the evolution of the IC pathway across plant species, highlighting the need for systematic evolutionary and functional analyses of the SA biosynthesis genes *ICS*, *EDS5*, and *PBS3*.

In this study, we aimed to elucidate the evolutionary origins of the IC pathway for SA biosynthesis through systematic phylogenetic analysis of *ICS*, *EDS5*, and *PBS3*, combined with functional reconstitution of the IC pathway in *Nicotiana benthamiana* and complementation of *A. thaliana* mutants using homologous genes of *ICS*, *EDS5*, and *PBS3* from diverse plant species. Our findings suggest that the IC pathway emerged within Brassicales.

## RESULTS

### The phylogenetic analysis of the IC pathway genes *ICS1*, *EDS5*, *PBS3*, and *EPS1*

To investigate the evolutionary origin and conservation of the IC pathway, we first identified homologous sequences of the IC pathway genes *ICS1*, *EDS5*, *PBS3*, and *EPS1* from *A. thaliana* across various plant species. Our reference point in this study is *A. thaliana*, which serves as the baseline for discussing the timing of divergence among various plant species. To ensure a comprehensive evolutionary framework, we included key taxa representing major plant lineages (Figure 1). These taxa included the liverwort *Marchantia polymorpha*, representing bryophytes, which is a sister group to tracheophytes (vascular plants), and the lycophytes *Selaginella moellendorffii*, which is a sister lineage to spermatophytes (gymnosperms and angiosperms). We also incorporated *Amborella trichopoda,* which is the sister species to all extant angiosperms. To assess Monocots, we included *Oryza sativa* (rice), *Zea mays* (maize), *Eleusine coracana* (finger millets), and *Thinopyrum intermedium*. For Magnoliids, we selected *Cinnamomum kanehirae* and *Liriodendron tulipifera*. To cover Eudicots, we encompassed representatives from Rosids and Asterids. Within Asterids, we included *Solanum tuberosum* (potato) and *Solanum lycopersicum* (tomato).

**Figure 1.**
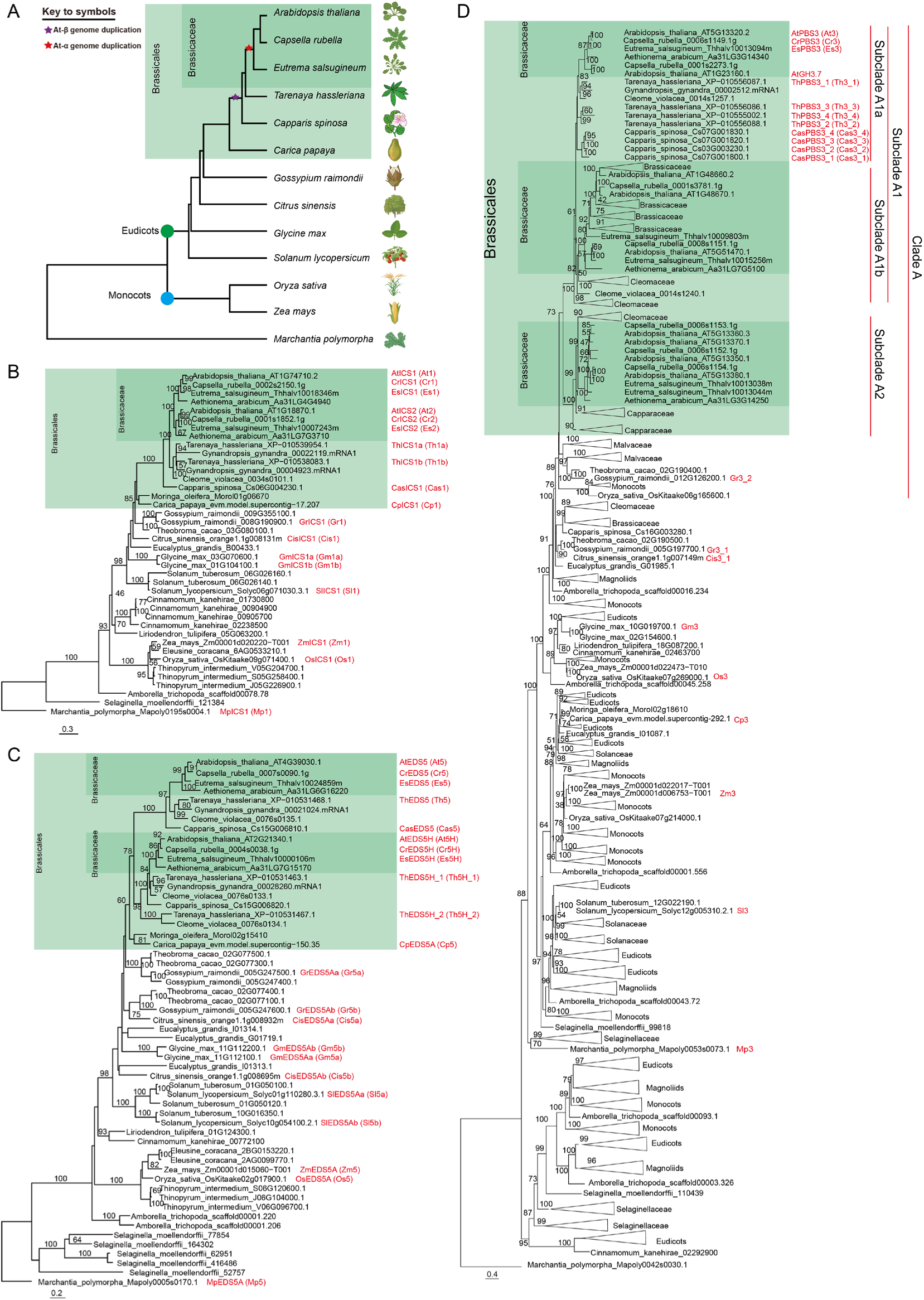
Phylogenic analysis of genes in the IC-based pathway across plant species. (**A**) Phylogenetic tree of the plant species analyzed in this study. (**B-D**) Phylogenic trees of ICS (**B**), EDS5 (**C**), and PBS3 (**D**) homologs from various plant species. Trees were generated based on full-length amino acid sequences using maximum likelihood method. Triangles represent collapsed branches. The scale bar denotes a phylogenetic distance of amino acid substitutions per site. Bootstrap values, derived from 1000 replications, are indicated at branch nodes. Some illustrations were created using BioRender. (For more detailed phylogenetic trees, see Datasets 1-3).

As *A. thaliana* belongs to the Brassicaceae family within the Brassicales order in Rosids, we focused extensively on species within Brassicales, including both Brassicaceae and non-Brassicaceae members, as well as species from other orders within Rosids. The Brassicaceae species examined, in order of phylogenetic proximity to *A. thaliana*, included *Capsella rubella*, *Eutrema salsugineum*, and *Aethionema arabicum*. The non-Brassicaceae Brassicales species analyzed, in order of phylogenetic proximity to *A. thaliana*, encompassed *Tarenaya hassleriana* and *Cleome violacea*, followed by *Capparis spinosa* (caper), with *C. papaya* (papaya) and *Moringa oleifera* representing early-diversifying lineages within Brassicales. To further refine the evolutionary context, we included taxa from closely related orders such as *Gossypium raimondii* (cotton) and *Theobroma cacao* from the order Malvales, which is the closest sister clade of Brassicales. We also selected *Citrus sinensis* (orange) from the order Sapindales, *Eucalyptus grandis* from the order Myrtales, which diversified before the divergence of Sapindales and after Fabales, and *Glycine max* (soybean) from the order Fabales. Within Brassicales, the At-β WGD event occurred after the divergence of *C. papaya* and *M. oleifera*, and the At-α WGD occurred in the common ancestor of Brassicaceae after the divergence of *T. hassleriana* and *C. violacea* lineages (39). This extensive taxon sampling allowed us to establish a robust phylogenetic framework to trace the evolutionary trajectory of the IC pathway genes across land plants.

To retrieve homologous sequences, perform sequence alignments, and construct phylogenetic trees, we utilized a previously developed pipeline (40). Phylogenetic analysis revealed that *ICS* genes are conserved across land plants (Figure 1B; see Dataset 1 for a more detailed phylogenetic tree). Gene duplication events were observed in several lineages. For instance, the *ICS* gene in the common ancestor of the Brassicaceae appears to have duplicated, as reported in *A. thaliana* (29), likely driven by the At-α WGD (41). This resulted in two distinct *ICS* clades, one containing *A. thaliana ICS1* and the other *ICS2*. While *A. thaliana ICS1* expression is inducible upon pathogen infection, *ICS2* is not (17). Consistently, we observed that *C. rubella* and *E. salsugineum ICS1* expression was upregulated following infection with the bacterial pathogen *Pseudomonas syringae* pv. *tomato* DC3000 (Supplementary Figure S1), suggesting that pathogen-inducible *ICS1* expression is a conserved trait within Brassicaceae.

While homologs of *EDS5* exist across land plants, the topology of the *EDS5* phylogenetic tree suggests that the ancestral gene of *EDS5* (*EDS5A*) duplicated after the divergence of *M. oleifera* and *C. papaya* within Brassicales (Figure 1C; see Dataset 2 for a more detailed phylogenetic tree). One of these duplicated lineages includes *A. thaliana EDS5*, while the other contains *A. thaliana EDS5H*, whose specific biological function remains unknown. Branch length analysis revealed that the *EDS5* clade has a longer branch than the *EDS5H* clade (Figure 1C), suggesting that greater evolutionary changes occurred in the *EDS5* clade, possibly associated with neofunctionalization and that genes in the *EDS5H* clade retained the ancestral function.

The GH3 gene family extensively diversified through many gene duplication and deletion events in various stages of the evolution of angiosperms (Figure 1D and Dataset 3), suggesting the requirement of highly diverse substrate specificities for the specialized GH3 functions in different plant lineages. The clade that contains the *A. thaliana* genes, At1g23160 (*AtGH3.7*), At1G48660, At1G48670, At5g13320 (*AtPBS3*), At5G13350, At5g13360, At5G13370, At5g13380, and At5g51470 diversified from the rest of the *GH3* gene family in the common ancestor of Eudicots and Monocots or earlier, evidenced by inclusion of the Monocot genes, such as *O. sativa* OsKitaake 06g165600. We named this clade Clade A. In the *AtPBS3* lineage within Clade A, one gene duplication event led to two subclades, one subclade (Subclade A1) containing At1g23160 (*AtGH3.7*), At1g48660, At1g48670, At5g13320 (*AtPBS3*), and At5g51470 and the other subclade (Subclade A2) containing At5g13350, At5g13360, At5g13370, and At5g13380. Subclade A1 was further diversified to Subclade A1a containing At5g13320 (*AtPBS3*) and At1g23160 (*AtGH3.7*) and Subclade A1b containing At1g48660, At1g48670, and At5g51470. Subclade A1a was further diversified around the time of the Brassicaceae family divergence into a subclade containing At5g13320 (*AtPBS3*) and the other subclade containing At1g23160 (*AtGH3.7*), which functions redundantly with *PBS3* in the IC pathway but with significantly weaker activity (42). Since the *A. thaliana* genes in Subclade A2 are very closely located on a chromosome to At5g13320 (*AtPBS3*) in Subclade A1a, it is likely that the diversification of Subclades A1 and A2 was initiated by a local gene duplication event and that a direct ancestor of At5g13320 (AtPBS3) was the MRCA of Subclade A1. Interestingly, Clade A does not include *C. papaya* and *M. oleifera* genes, while species of Malvales (*G. raimondii* and *T. cacao*), which is the sister order of Brassicales, have Clade A genes. Since the gene deletion is observed in both of the sister families, Caricaceae and Moringaceae, it is unlikely that an apparent deletion was a mistake in genome sequencing and assembly of *C. papaya* and *M. oleifera*. With a parsimonious inference that only requires a single gene deletion event, the gene duplication for Subclade A1 and A2 occurred after the divergence of *C. papaya* and *M. oleifera* and before the divergence of *C. spinosa* (evidenced by the inclusion of *C. spinosa* genes in Subclades A1 and A2) from the *A. thaliana* lineage, and the deletion of the Clade A gene occurred in *C. papaya* and *M. oleifera* lineages after the divergence from the *A. thaliana* lineage before the divergence between *C. papaya* and *M. oleifera*. With the inferences that diversification of Subclade A1 and A2 occurred after diversification of the *C. papaya* and *M. oleifera* lineages from the *A. thaliana* lineage and the MRCA of Subclade A1 is the direct ancestor of *AtPBS3*, we could define the MRCA of the Subclade A1 *C. spinosa* genes *Cas3_1*, *Cas3_2*, *Cas3_3*, and *Cas3_4* and *T. hassleriana* genes *Th3_1*, *Th3_2*, *Th3_3*, and *Th3_4* as the ortholog of *AtPBS3*, but we could not define the *AtPBS3* orthologs in the species diversified at or earlier than the diversification of the *C. papaya* and *M. oleifera* lineage from the *A. thaliana* lineage.

Consistent with a previous report (18), phylogenetic analysis confirmed that the *A. thaliana EPS1* clade is restricted to Brassicaceae (Supplementary Figure S2).

### Functional conservation of Brassicaceae IC pathway genes in SA biosynthesis

To assess the functional conservation of the IC pathway genes in SA biosynthesis, we employed a previously established assay in *Nicotiana benthamiana* (18). In this system, increased SA accumulation occurs only when functional *ICS1*, *EDS5*, and *PBS3* are co-expressed (Figure 2A). Consistent with the previous report (18), we observed significant accumulation of SA and SAG when *A. thaliana ICS1*, *EDS5*, and *PBS3* were co-expressed, whereas no substantial accumulation was detected when only two of these genes were expressed (Figure 2B and C). Similarly, co-expression of *ICS1*, *EDS5*, and *PBS3* from *C. rubella* and *E. salsugineum* resulted in comparable SA and SAG accumulation to *A. thaliana* genes (Figure 2B and C), suggesting that the functions of ICS1, EDS5, and PBS3 are conserved in Brassicaceae.

**Figure 2.**
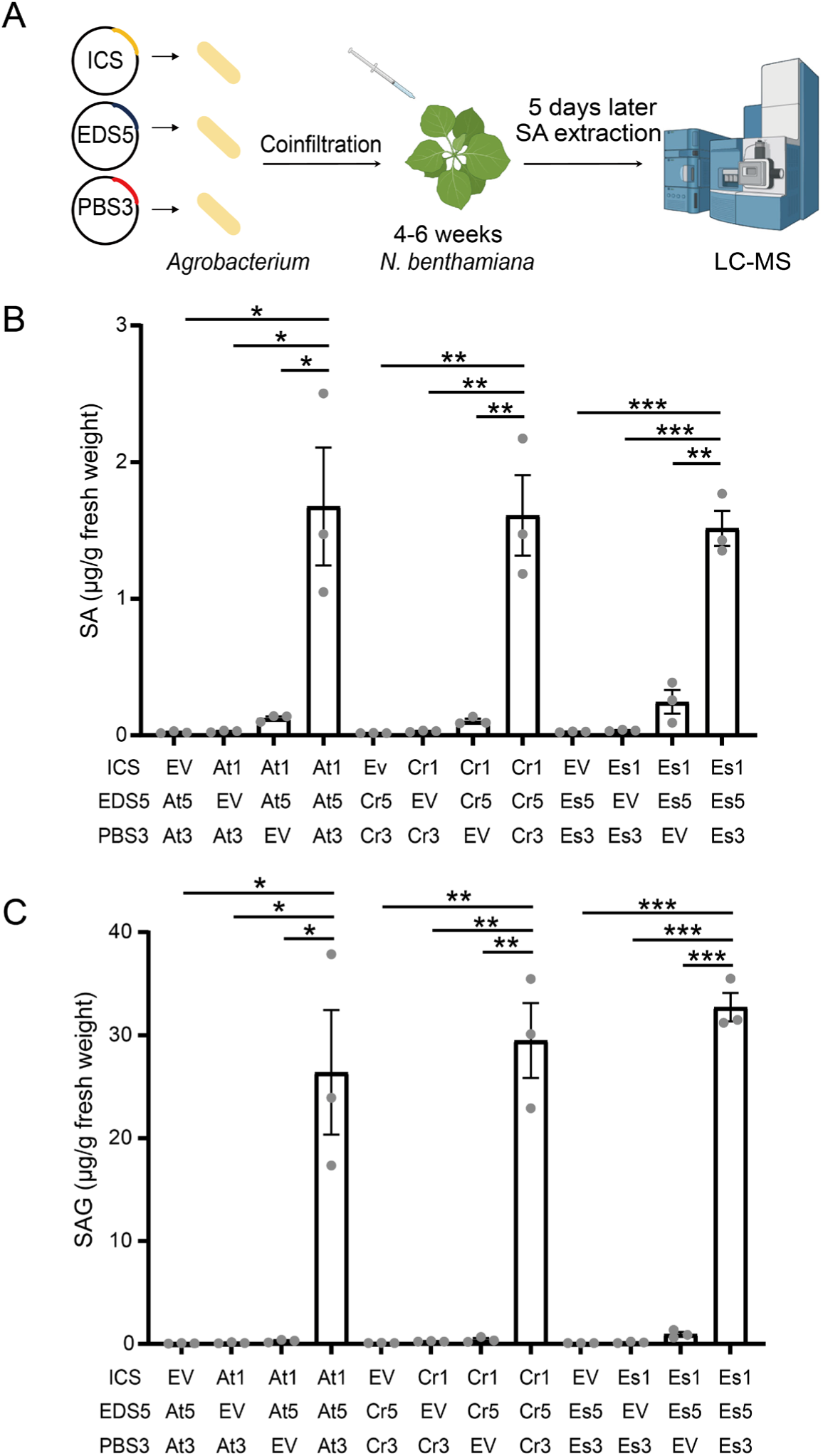
Brassicaceae ICS1, EDS5, and PBS3 are functional for SA biosynthesis in *N. benthamiana*. (**A**) Schematic representation of the experimental design. (**B** and **C**) *Agrobacterium* strains carrying *ICS-Myc*, *EDS5-YFP*, or *PBS3-YFP* from the *Brassicaceae* species (*A. thaliana*, *C. rubella*, and *E. salsugineum*) were co-infiltrated into leaves of 4 to 6-week-old *N. benthamiana* plants. SA and SAG levels in leaf extracts were quantified at 5 days post-inoculation (dpi). EV represents the empty vector control. Data represent means ± SEM from 3 biological replicates. Asterisks indicate significant differences (\**P <* 0.05, \*\**P <* 0.01, and \*\*\**P* < 0.001, two-tailed Student’s t-tests). Some illustrations were created using BioRender.

We noticed that *N. benthamiana* appears to have a weak PBS3-like activity (Figure 2B). As IC can be non-enzymatically converted to SA (43), it might be possible that this low level of SA accumulation represents non-enzymatically converted SA when a large amount of IC is produced. Alternatively, some GH3 enzymes in *N. benthamiana* may happen to have IC-converting activity. Note that when IC is converted to SA non-enzymatically, such conversion-enhancing enzyme may not have the glutamate-conjugating activity like PBS3 but have a different enzyme activity.

### ICSs from Brassicales exhibit strong activity in SA biosynthesis

To investigate the functional conservation of *ICS* genes across diverse plant species, we expressed *ICS* genes from different species together with *A. thaliana EDS5* and *PBS3* in *N. benthamiana*. The results showed that *ICS1* and *ICS2* from the Brassicaceae species *A. thaliana*, *C. rubella*, and *E. salsugineum* significantly increased SA accumulation. In contrast, *ICS* genes from non-Brassicales species, including *G. max*, *S. lycopersicum*, *Z. mays*, *O. sativa*, and *M. polymorpha*, led to little or no increase in SA accumulation (Figure 3A and B). These findings suggest that ICSs in Brassicaceae have evolved higher SA biosynthesis activity than those in non-Brassicales plant lineages.

**Figure 3.**
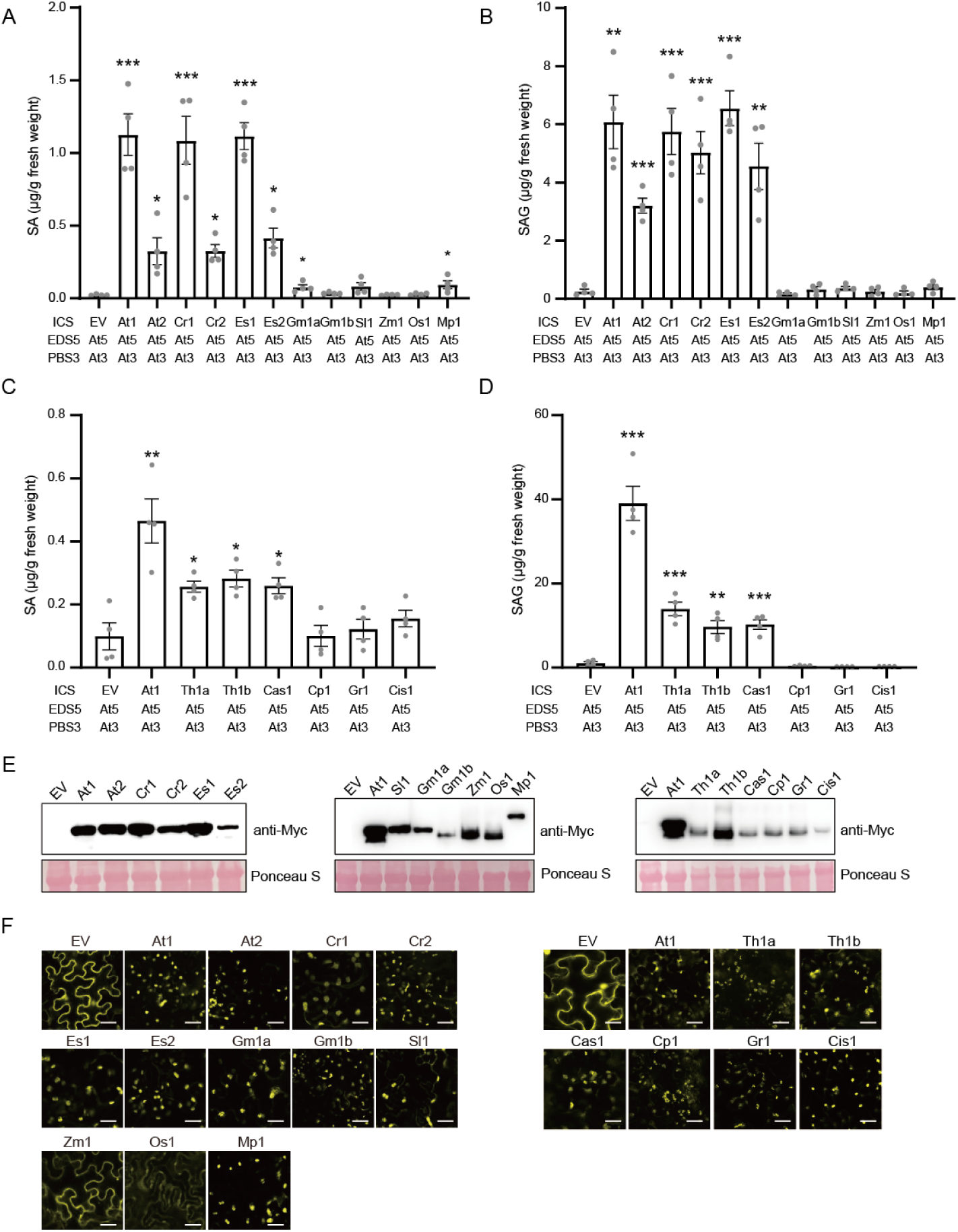
Brassicales ICS homologs except for *C. papaya* are functional for SA biosynthesis in *N. benthamiana*. *Agrobacterium* strains carrying *ICS-Myc* from various species were co-infiltrated with *Agrobacterium* strains carrying *A. thaliana EDS5-YFP* and *PBS3-YFP* into leaves of 4 to 6-week-old *N. benthamiana* plants. (**A-D**) SA and SAG levels in leaf extracts were quantified at 5 dpi. EV represents the empty vector control. Data represent means ± SEM from 4 biological replicates. Asterisks indicate significant differences compared to EV (\**P <* 0.05, \*\**P <* 0.01, and \*\*\**P* < 0.001, two-tailed Student’s t-tests). (**E**) *Agrobacterium* strains carrying *ICS-Myc* were infiltrated into *N. benthamiana* leaves. Total protein was extracted from the infiltrated leaves at 2 dpi, and ICS protein expression was assessed by immunoblotting using an anti-Myc antibody. (**F**) *Agrobacterium* strains carrying *ICS-YFP* were infiltrated into *N. benthamiana* leaves. ICS subcellular localization in epidermal cells expressing ICS-YFP was analyzed by confocal microscopy at 2 dpi. Scale bars, 20 µm.

To further assess the functional conservation of *ICS*, we examined the non-Brassicaceae Brassicales species *T. hassleriana*, *C. spinosa*, and *C. papaya* as well as species from closely related sister lineages, *G. raimondii* and *C. sinensis*. Reconstitution of the IC pathway in *N. benthamiana* revealed the expression of *ICS* genes from *T. hassleriana* and *C. spinosa* resulted in elevated SA and SAG accumulation (Figure 3C and D). However, *ICS* genes from *C. papaya*, *G. raimondii*, and *C. sinensis* did not (Figure 3C and D). Immunoblotting assays confirmed the expression of all ICS-Myc proteins in *N. benthamiana* (Figure 3E).

To determine whether differences in ICS activity were associated with subcellular localization, we performed confocal microscopy analysis of ICS-YFP fusion proteins. Most ICS-YFP proteins localized to chloroplasts (Figure 3F and Supplementary Figure S3), consistent with the previous report for *A. thaliana* ICS1 (29). However, ICS from Monocots (*Z. mays* and *O. sativa*) were predominantly localized to the cytosol (Figure 3G and Supplementary Figure S3). This cytosolic localization may explain why Monocot ICSs did not function as efficiently as *A. thaliana* ICS1. Alternatively, the chloroplast-localization peptide sequences in Monocot ICSs might be non-functional in *N. benthamiana* as a previous study reported that *O. sativa* ICS1 localizes to chloroplasts in rice protoplasts (26). Nevertheless, the fact that *C. papaya* ICS and other non-functional ICSs were chloroplast-localized indicates that chloroplast localization alone is not sufficient for enhanced SA biosynthesis activity in this system. Together, these results indicate that while some species, such as *M. polymorpha*, exhibit weak ICS activity, strong SA biosynthesis activity is observed only in Brassicales species that diversified after the divergence of *M. oleifera* and *C. papaya* lineages. To further confirm their functionality in the IC pathway, we performed complementation assays using *A. thaliana ics1* (*sid2*) mutants. Consistent with our observations in *N. benthamiana* reconstitution assays, *ICS1* genes from the Brassicaceae *A. thaliana*, *C. rubella*, and *E. salsugineum*, as well as the non-Brassicaceae Brassicales *T. hassleriana* and *C. spinosa*, successfully complemented the *sid2* mutant, restoring pathogen-inducible SA and SAG accumulation (Supplementary Figure S4). In contrast, *ICS* from other species, including the Brassicales *C. papaya*, failed to complement the *ics1* mutant phenotype (Supplementary Figure S4). These results suggest that ICS activity for SA biosynthesis was significantly enhanced within Brassicales species that diversified after the divergence of *M. oleifera* and *C. papaya* lineages.

### Enhanced salt bridge network may contribute to the high ICS activity in Brassicales

To understand why ICS enzymes from certain Brassicales species exhibited enhanced activity for SA biosynthesis, we performed structural predictions of *A. thaliana* ICS1 using AlphaFold2 and compared the results with previously determined ICS structures (44). Our analysis revealed that E366 in AtICS1 corresponds to E197 in *E. coli* ICS, a residue essential for ICS enzymatic function. In *E. coli*, the carboxyl group of E197 is oriented toward the C4 position of the bound IC and functions as the general acid for the loss of the C4 hydroxyl group from IC (44). Consistent with this and a recent publication of *A. thaliana* ICS1 structure (45), a mutated version of *A. thaliana* ICS (AtICS1^E366A^) failed to induce SA accumulation in *N. benthamiana* reconstitution assays, despite exhibiting proper chloroplast localization (Supplementary Figures S5A-C and S6). A key structural feature distinguishing *A. thaliana* ICS1 from bacterial ICS is an extensive salt bridge network that stabilizes E366. This network comprises residues R130, D132, R135, D346, R367, E379, R438, and E509 (Supplementary Figure S5D). Among these, D132, R135, D346, and R367 are predominantly conserved in Brassicales ICSs that retained strong SA biosynthesis function in *N. benthamiana* reconstitution assays (Supplementary Figure S7). To assess the contribution of these residues to ICS1 activity, we introduced alanine mutations at these positions. Our results indicate that while D132A mutation had a minimal effect on AtICS1 activity, R135A, D346A, and R367A mutations substantially reduced AtICS1 activity, without affecting subcellular localization (Supplementary Figure S5A-C and S6). To further dissect their role, we generated double and triple mutants (AtICS1^D132AR135A^, AtICS1^R135AD346A^, AtICS1^R135AR367A^, AtICS1^R346AR367A^, and AtICS1^R135AD346AR367A^). While AtICS1^D132AR135A^ and AtICS1^D346AR367A^ retained weak SA biosynthesis activity, AtICS1^R135AD346A^, AtICS1^R135AR367A^, and AtICS1^R135AD346AR367A^ completely lost their functions, despite proper subcellular localization (Supplementary Figures S5A-C and S6). Immunoblotting assays confirmed the expression of all ICS-Myc proteins (Supplementary Figure S5E). These findings suggest that the enhanced salt bridge network of the ICSs in Brassicales, except for *C. papaya*, plays a crucial role in stabilizing key catalytic residues, thereby contributing to their enhanced activity.

### PBS3 function is restricted to Brassicales species that diversified after the divergence of *M. oleifera* and *C. papaya*

Our phylogenetic analysis suggests that PBS3 function might be confined to Brassicales species that diversified after the divergence of *M. oleifera* and *C. papaya* lineages (Figure 1D). To experimentally validate this hypothesis, we conducted *N. benthamiana* reconstitution assays. Co-expression of *PBS3* from *C. rubella* and *E. salsugineum* with *A. thaliana ICS1* and *EDS5* significantly induced SA and SAG accumulation (Figure 4A and B), suggesting that PBS3 function is conserved in Brassicaceae. In contrast, the closest homologs of *PBS3* from the non-Brassicales species, including *S. lycopersicum*, *G. max*, *Z. mays*, *O. sativa*, *M. polymorpha*, *C. sinensis*, and *G. raimondii*, as well as the Brassicales *C. papaya*, failed (Figure 4A-D). Notably, some *PBS3* homologs from *T. hassleriana* and *C. spinosa*, which belong to Subclade A1a containing *A. thaliana PBS3* and GH3.7, retained SA biosynthesis activity. Subcellular localization assays in *N. benthamiana* demonstrated that all PBS3-YFP fusion proteins localized to the cytosol, and immunoblotting confirmed their expression (Figure 4E and F and Supplementary Figure S8).

**Figure 4.**
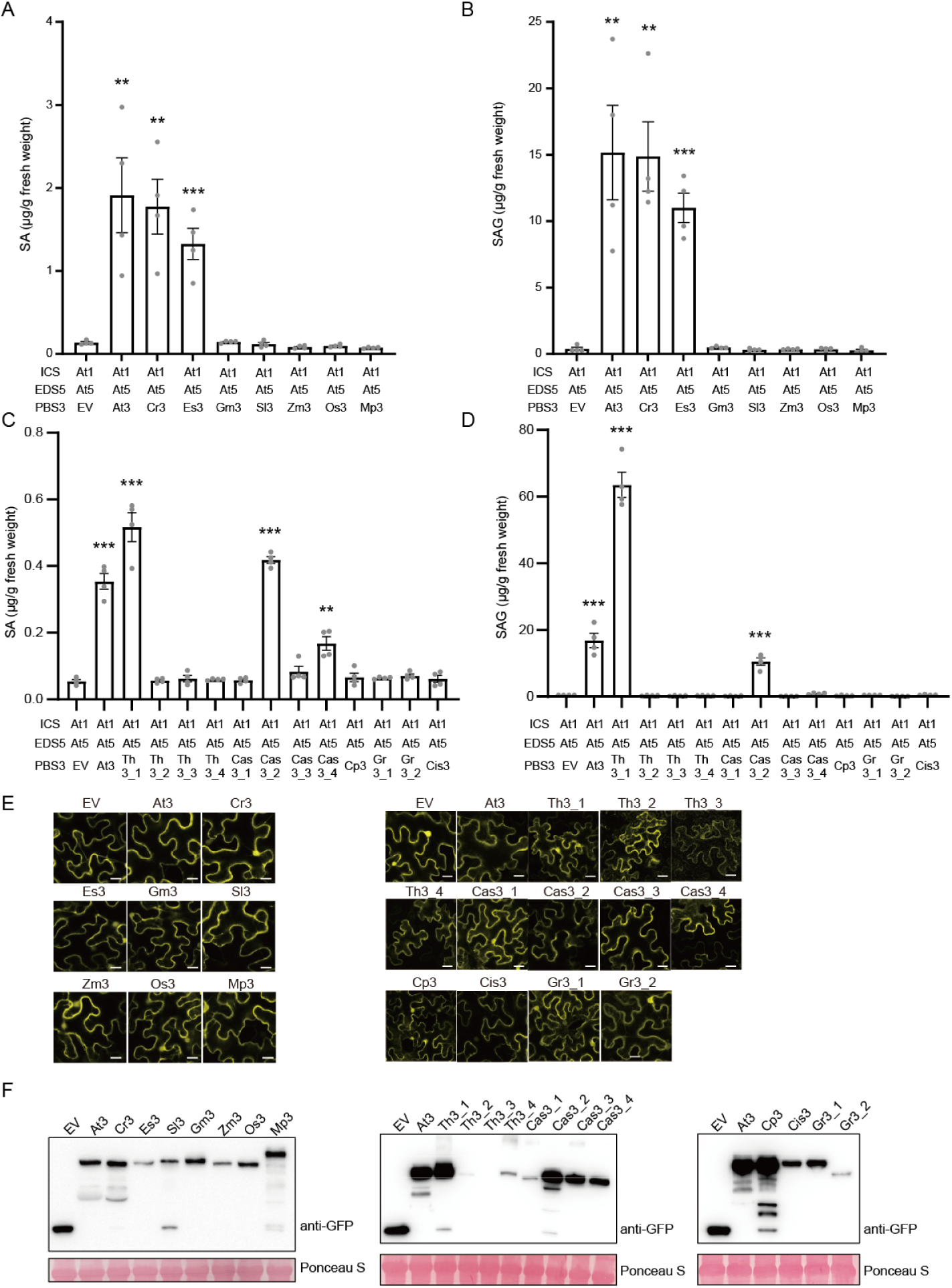
Brassicales PBS3 homologs, except for *C. papaya*, are functional for SA biosynthesis in *N. benthamiana*. *Agrobacterium* strains carrying *PBS3-YFP* from various species were co-infiltrated with *Agrobacterium* strains carrying *A. thaliana ICS-Myc* and *EDS5-YFP* into leaves of 4 to 6-week-old *N. benthamiana* plants. (**A-D**) SA and SAG levels in leaf extracts were quantified at 5 dpi. EV represents the empty vector control. Data represent means ± SEM from 4 biological replicates. Asterisks indicate significant differences compared to EV (\**P <* 0.05, \*\**P <* 0.01, and \*\*\**P* < 0.001, two-tailed Student’s t-tests). (**E**) *Agrobacterium* strains carrying *PBS3-YFP* were infiltrated into *N. benthamiana* leaves. PBS3 subcellular localization in epidermal cells expressing PBS3-YFP was analyzed by confocal microscopy at 2 dpi. Scale bars, 20 µm. (**F**) *Agrobacterium* strains carrying *PBS3-YFP* were infiltrated into *N. benthamiana* leaves. Total protein was extracted from the infiltrated leaves at 2 dpi, and PBS3 protein expression was assessed by immunoblotting using an anti-GFP antibody.

Given that PBS3 belongs to the large GH3 protein family, it is possible that other GH3 members function as PBS3. To explore this possibility, we examined the functions of other *A. thaliana* GH3 family members in *N. benthamiana* reconstitution assays. The results revealed that only PBS3 (GH3.12) and GH3.7, which belong to Subclade A1a, exhibited SA biosynthesis activity (Figure 1D and Supplementary Figure S9A and B). This was observed despite proper cytosolic localization and expression in all tested GH3 proteins, except for GH3.8, which was not detected in immunoblotting (Supplementary Figures S9A and B and S10). Together with our phylogenetic analysis, these results suggest that the PBS3 function in the IC pathway is restricted to Subclade A1a in Brassicales species that diversified after the divergence of *M. oleifera* and *C. papaya* lineages.

### Neofunctionalization of EDS5 within Brassicales

The longer branch length in the *A. thaliana EDS5* clade compared to the *A. thaliana EDS5H* clade following gene duplication (Figure 1C) suggests that the *EDS5* lineage underwent neofunctionalization, leading to the emergence of a role for *EDS5* in the IC pathway. To test this hypothesis, we performed *N. benthamiana* reconstitution assays. The results revealed that co-expression of *EDS5* from *A. thaliana*, *C. rubella*, and *E. salsugineum* with *A. thaliana ICS1* and *PBS3* led to strong SA and SAG accumulation (Figure 5A and B). In contrast, *EDS5H* from these species, as well as homologs of *EDS5* (*EDS5A*) from non-Brassicales species and the Brassicales species *C. papaya*, failed (Figure 5A-D). Notably, while *EDS5* from the non-Brassicaceae Brassicales species *T. hassleriana* and *C. spinosa* successfully induced SA and SAG accumulation, *EDS5H* from these species did not (Figure 5C and D). These results were observed despite proper chloroplast membrane localization and protein expression of all tested EDS5-related proteins (Figure 5E and F and Supplementary Figure S11). Consistent with these findings, *EDS5* from the Brassicaceae species *A. thaliana*, *C. rubella*, and *E. salsugineum*, as well as the non-Brassicaceae Brassicales species *T. hassleriana* and *C. spinosa*, successfully complemented the *eds5* mutant, restoring pathogen-inducible SA and SAG accumulation (Supplementary Figure S12). In contrast, *EDS5A* from other species, including *C. papaya*, failed to complement the *eds5* phenotype. These findings strongly support the hypothesis that *EDS5* underwent neofunctionalization following gene duplication to participate in the IC pathway for SA biosynthesis within Brassicales.

**Figure 5.**
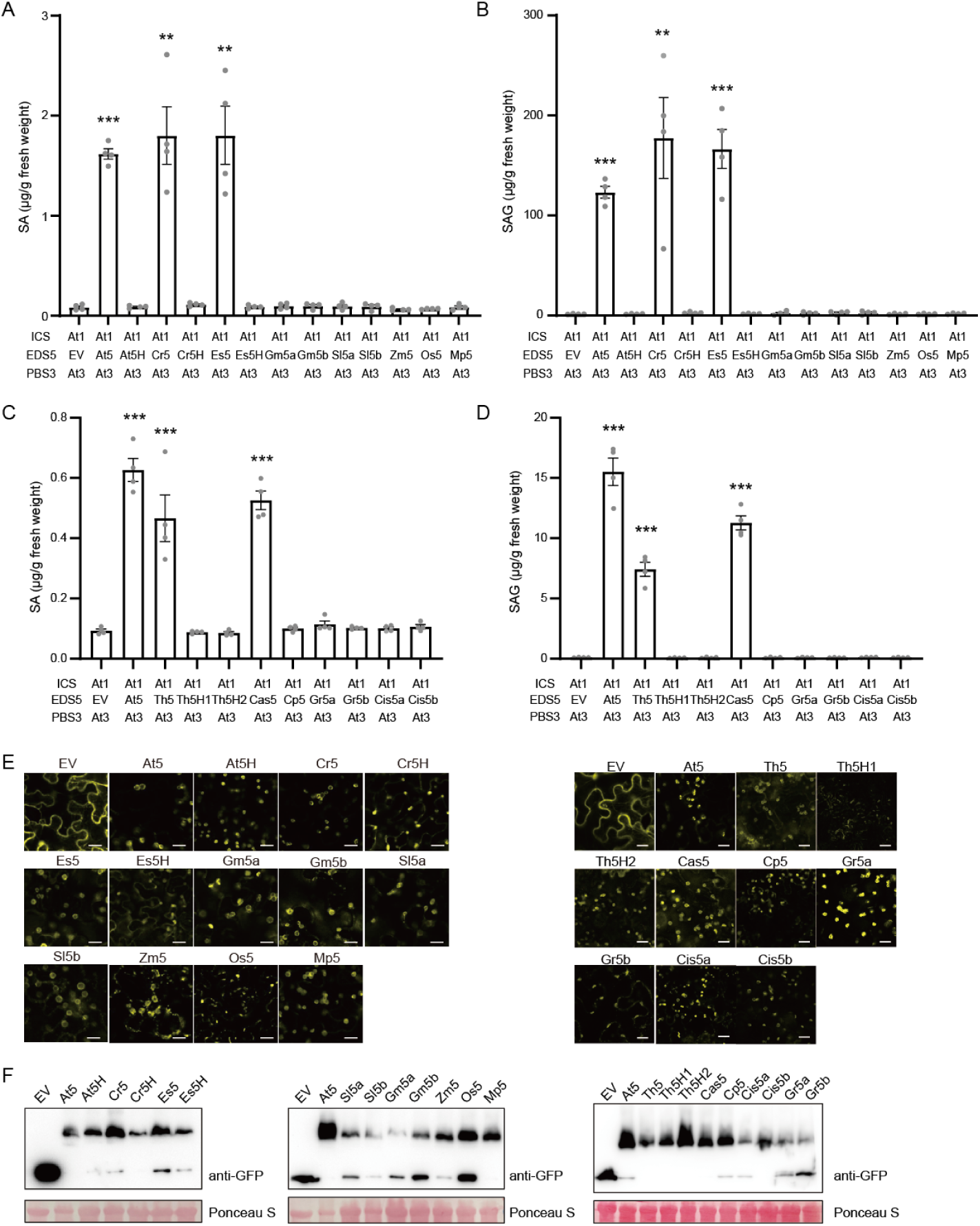
Brassicales EDS5 homologs except for *C. papaya* are functional for SA biosynthesis in *N. benthamiana*. *Agrobacterium* strains carrying *EDS5-YFP* from various species were co-infiltrated with *Agrobacterium* strains carrying *A. thaliana ICS-Myc* and *PBS3-YFP* into leaves of 4 to 6-week-old *N. benthamiana* plants. (**A-D**) SA and SAG levels in leaf extracts were quantified at 5 dpi. EV represents the empty vector control. Data represent means ± SEM from 4 biological replicates. Asterisks indicate significant differences compared to EV (\**P <* 0.05, \*\**P <* 0.01, and \*\*\**P* < 0.001, two-tailed Student’s t-tests). (**E**) *Agrobacterium* strains carrying *EDS5-YFP* were infiltrated into *N. benthamiana* leaves. EDS5 subcellular localization in epidermal cells expressing *EDS5-YFP* was analyzed by confocal microscopy at 2 dpi. Scale bars, 20 µm. (**F**) *Agrobacterium* strains carrying *EDS5-YFP* were infiltrated into *N. benthamiana* leaves. Total protein was extracted from the infiltrated leaves at 2 dpi, and EDS5 protein expression was assessed by immunoblotting using an anti-GFP antibody.

At present, whole genome sequence data for non-Brassicaceae Brassicales species beyond those used in this study are not available. Nevertheless, to further refine the evolutionary trajectory of *EDS5* within Brassicales, we selected the additional Brassicales species—*Reseda odorata*, *Batis maritima*, and *Limnanthes douglasii*—which diversified after the divergence of *C. papaya* but before *C. spinosa*, in order of phylogenetic proximity to *A. thaliana* (Supplementary Figure S13). We cultivated these species, extracted RNA, and performed RNA sequencing followed by *de novo* transcriptome assembly. Homologous sequences to *A. thaliana EDS5* were identified and tested in *N. benthamiana* reconstitution assays. The results showed that only *R. odorata EDS5* induced SA and SAG accumulation, whereas *EDS5A* genes from *B. maritima* and *L. douglasii* did not, despite all EDS5-related proteins localizing to chloroplast membrane and being expressed as confirmed by immunoblotting (Supplementary Figure S13). These findings suggest that the duplication of the ancestral gene of *EDS5*, followed by the *EDS5* neofunctionalization, occurred during Brassicales evolution within the time frame between the divergence of *B. maritima* and *R. odorata*.

### Arg and Asn residues in the C-lobe of EDS5 are essential for the function in SA biosynthesis

MATE transporter family proteins typically contain two substrate binding sites, located in the N-lobe and C-lobe (46). Since EDS5 is hypothesized to transport IC from chloroplasts to the cytosol, we predicted the structure of *A. thaliana* EDS5 using AlphaFold2 to determine whether it harbors potential substrate-binding sites. The structural predictions combined with amino acid sequence alignment revealed putative key residues in the N-lobe (S293) and C-lobe (R366, N369, and Y516) that are almost exclusively conserved in EDS5 from Brassicales species with retained SA biosynthetic function in *N. benthamiana* reconstitution assays (Figure 6A and B). To assess the functional significance of these residues, we introduced alanine mutations at these positions and conducted *N. benthamiana* reconstitution assays. While S293A and Y516A mutations had no effect on *A. thaliana* EDS5 function in SA biosynthesis, single (R366A or N369A) and double (R366A N369A) mutations completely abolished SA biosynthetic activity (Figure 6C and D). This loss of function occurred despite proper chloroplast membrane localization and protein expression, as confirmed by fluorescent microscopy and immunoblotting (Figure 6E and F and Supplementary Figure S14). Consistent with these findings, the corresponding mutant versions of EDS5 from *C. rubella*, *E. salsugineum, T. hassleriana*, and *C. spinosa* are all non-functional in SA biosynthesis in *N. benthamiana* reconstitution assays (Supplementary Figures S15). Strikingly, *R. odorata* EDS5, which retained SA biosynthesis activity, possesses Arg and Asn residues at the corresponding positions, whereas EDS5As of *B. maritima* and *L. douglasii*, which were non-functional, lack these residues (Figure 6B). These findings strongly suggest that R366 and N369 in the EDS5 C-lobe are essential for its role in SA biosynthesis and are exclusively conserved in functional Brassicales EDS5. The evolution of these two amino acids may have been a key event in EDS5 neofunctionalization, and their presence could serve as a molecular marker for identifying functional EDS5.

**Figure 6.**
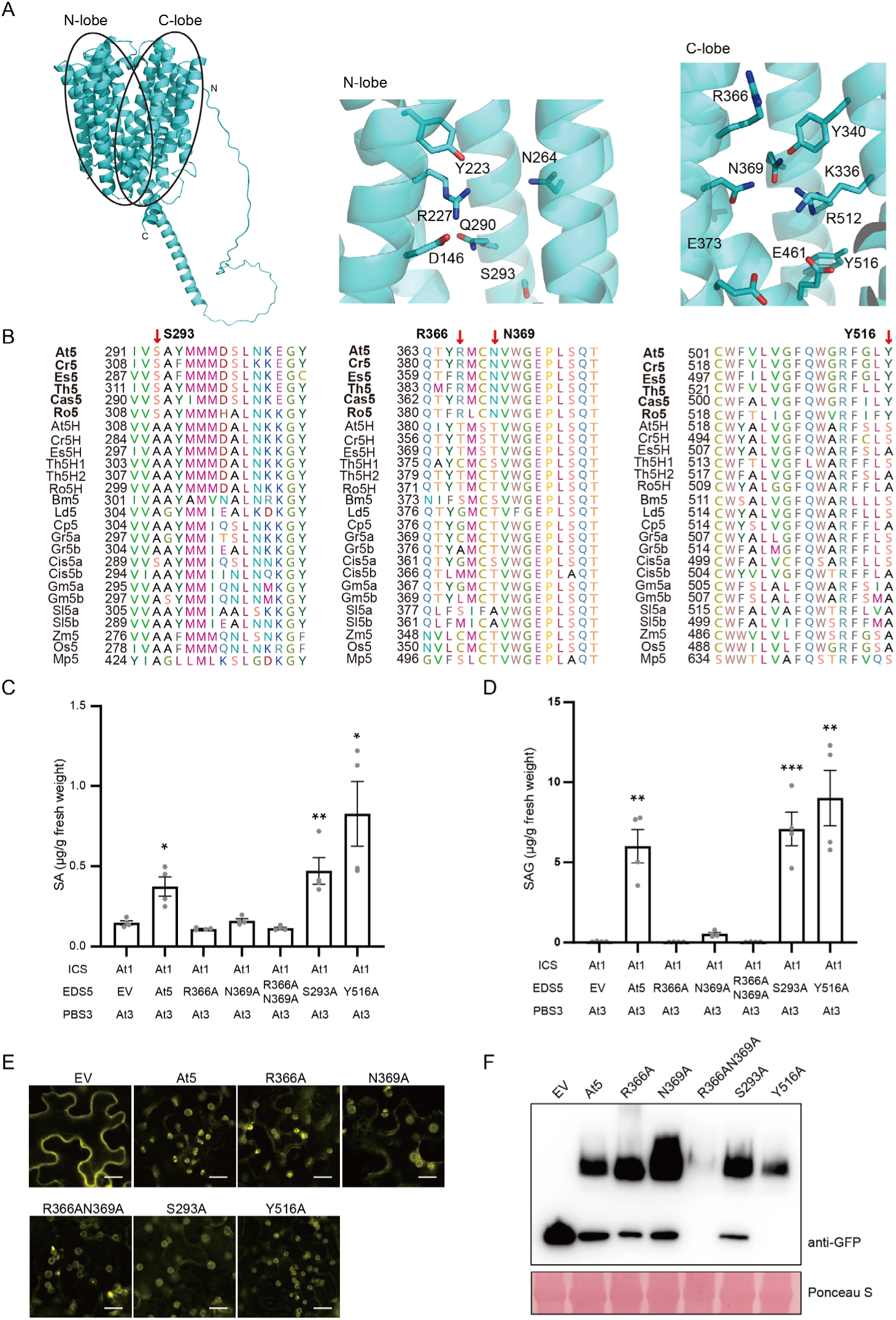
AtEDS5 residues R366 and N369 in the predicted IC binding site are essential for SA biosynthesis function. (**A**) Structural model of AtEDS5 predicted by AlphaFold2. Both the overall structure and close-up views of the candidate regions for the IC binding pockets (N-lobe and C-lobe) are shown. (**B**) Alignment of EDS5 homolog amino acid sequences from the plant species analyzed in this study. The red arrows indicate the mutated sites in EDS5. EDS5 homologs with bold letters are functional for SA biosynthesis in this study. (**C** and **D**) *Agrobacterium* strains carrying *ICS-Myc*, *EDS5-YFP* (or mutated *EDS5-YFP*), and *PBS3-YFP* from *A. thaliana* were co-infiltrated into leaves of 4 to 6-week-old *N. benthamiana* plants. SA and SAG levels in leaf extracts were quantified at 5 dpi. EV represents the empty vector control. Data represent means ± SEM from 4 biological replicates. Asterisks indicate significant differences compared to EV (\**P <* 0.05, \*\**P <* 0.01, and \*\*\**P* < 0.001, two-tailed Student’s t-tests). (**E**) *Agrobacterium* strains carrying *EDS5-YFP* were infiltrated into *N. benthamiana* leaves. EDS5 subcellular localization in epidermal cells expressing EDS5-YFP was analyzed by confocal microscopy at 2 dpi. Scale bars, 20 µm. (**F**) *Agrobacterium* strains carrying *EDS5-YFP* were infiltrated into *N. benthamiana* leaves. Total protein was extracted from the infiltrated leaves at 2 dpi, and EDS5 protein expression was assessed by immunoblotting using an anti-GFP antibody.

### *EDS5A* is not necessary for SA accumulation in soybean, maize, and rice

Our findings suggest that homologs of *EDS5* (*EDS5A*) do not play a major role in SA biosynthesis in non-Brassicales species. To further test this hypothesis, we analyzed *eds5A* mutants in *G. max*, *Z. mays*, and *O. sativa*. A *Z. mays* mutant was identified from an EMS-mutagenized maize B73 population, while *G. max* and *O. sativa* mutants were generated using CRISPR-Cas (Supplementary Figure S16). As *G. max* possesses two *EDS5A* homologs (*GmEDS5Aa* and *GmEDS5Ab*), we generated both a single *Gmeds5Ab* and the double *Gmeds5Aa/b* mutants. While we attempted to generate CRISPR *eds5A* mutants in *S. lycopersicum*, we were unable to obtain homozygous mutant plants from heterozygous mutant plants carrying mutations in the *S. lycopersicum EDS5A* homolog, likely due to its essential function in development. In *G. max*, *Gmeds5Ab* and *Gmeds5Aa/b* mutant plants retained the ability to accumulate large amounts of SA in response to infection by the bacterial pathogen *P. syringae* pv. *glycinea* Neau001 (Figure 7A and B). Similarly, in *Z. mays* and *O. sativa*, where SA accumulation is not pathogen-inducible, mutant plants maintained wild-type-like basal SA levels (Figure 7C-F). Consistent with these findings and in contrast to Brassicaceae species (Supplementary Figure S1), pathogen infection did not induce the expression of *ICSs* and *EDS5As* in *S. lycopersicum*, *G. max*, *Z. mays*, and *O. sativa*, despite the strong induction of the immune marker pathogenesis-related (PR) genes (Supplementary Figure S17). Taken together, these results further support the conclusion that the IC pathway is functionally restricted to specific Brassicales species.

**Figure 7.**
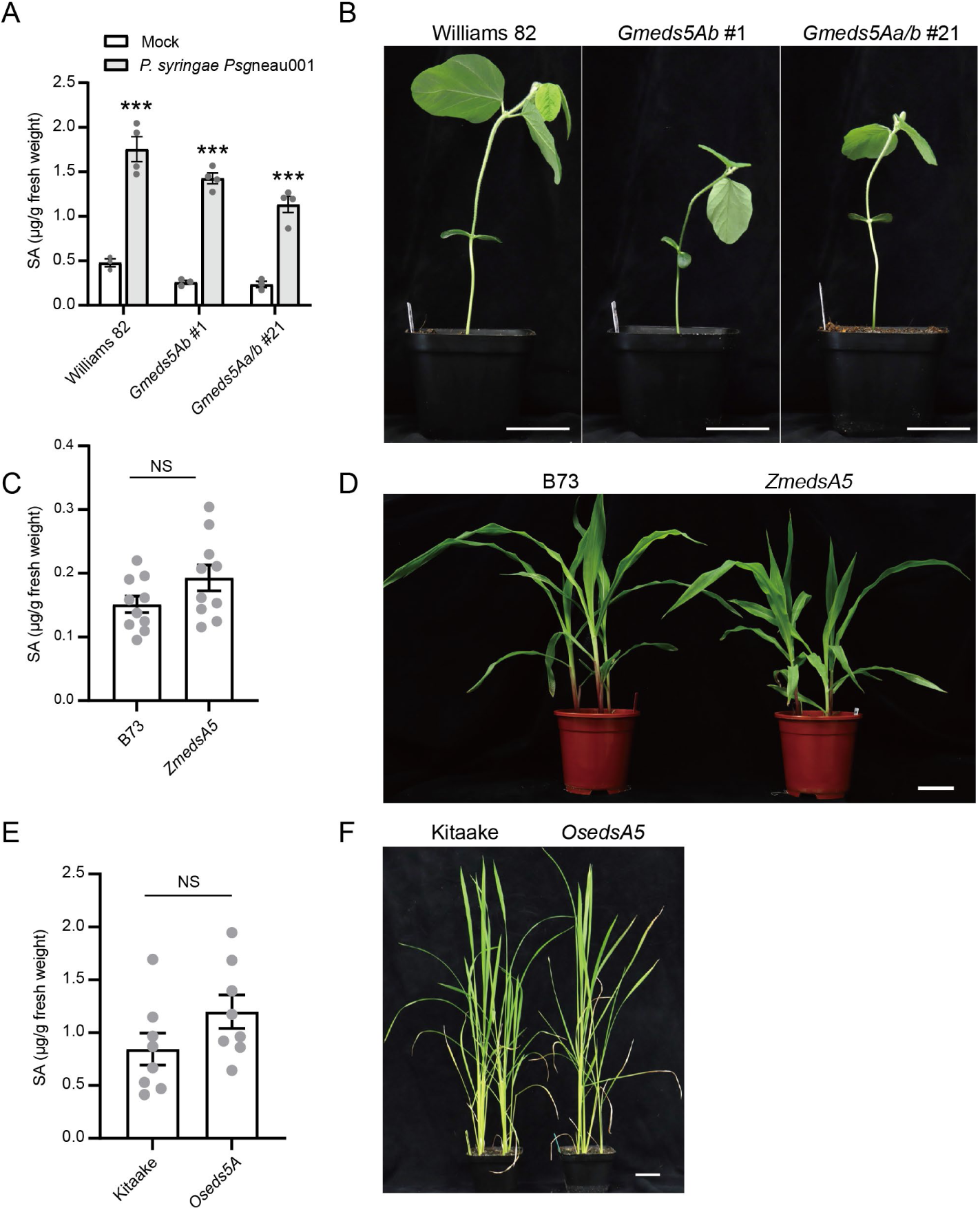
*EDS5A* is not essential for SA accumulation in soybean, maize, and rice. (**A**) Leaves of 2-week-old wild-type (Williams 82) and *Gmeds5Ab* and *Gmeds5Aa/b* soybean mutant plants were infected with *P. syringae Psg*neau001 (OD_600_=0.5) or mock. SA levels in leaf extracts were measured at 2 dpi. Data represent means ± SEM from at least 3 biological replicates. (**B**) Representative images of 2-week-old soybean plants. Scale bar=5 cm. (**C**) Basal SA levels in leaves of 3-week-old wild-type (B73) and *Zmeds5A* mutant maize plants. Data represent means ± SEM from 10 biological replicates. (**D**) Representative images of 3 -week-old maize plants. Scale bar=5 cm. (**E**) Basal SA levels in leaves of 4-week-old wild-type (Kitaake) and *Oseds5A* mutant rice plants. Data represent means ± SEM from 8 biological replicates. (**F**) Representative images of 4-week-old rice plants. Scale bar=1 cm. (**A, C, E**) Asterisks indicate significant differences between mock and pathogen treatments (\**P <* 0.05, \*\**P <* 0.01, and \*\*\**P* < 0.001, two-tailed Student’s t-tests). “NS” indicates no significant difference (*P* > 0.05).

## DISCUSSION

Our comprehensive phylogenetic analyses, integrated with functional assays in *N. benthamiana* and *A. thaliana* for core IC pathway genes—*ICS*, *EDS5*, and *PBS3*— alongside genetic studies of *EDS5A* in soybean, maize, and rice, provide strong evidence that the IC pathway is conserved exclusively in Brassicales species that diversified after the divergence of *M. oleifera* and *C. papaya* lineages. For EDS5, we further delineate that its neofunctionalization occurred during Brassicales evolution within the timeframe between the divergence of *B. maritima* and *R. odorata*. The absence of duplicated *EDS5* and *EDS5H* homologs in *B. maritima* and *L. douglasii*, despite their post-At-β WGD diversification, and the presence of only *EDS5A*, which was non-functional, suggests that a localized gene duplication rather than the At-β WGD contributed to the duplication of *EDS5A*. Further, our study and a previous report (18) suggest that the subsequent Brassicaceae-specific innovation of *EPS1* may have further enhanced the efficiency of IC-based SA biosynthesis within Brassicaceae.

Our findings indicate that the evolution of the IC pathway necessitated at least three key adaptations: (1) enhanced ICS activity, (2) neofunctionalization of *EDS5*, and (3) the evolution of a specialized *PBS3*. Enhanced ICS activity is associated with the strengthening of a salt bridge network around a key catalytic amino acid, thereby improving enzymatic efficiency. Neofunctionalization of EDS5 following the gene duplication is linked to amino acid substitutions in the substrate-binding site of the C-lobe, which likely refined its transport specificity. The *PBS3* gene clade is uniquely found in a Brassicales clade that diversified after the divergence of *M. oleifera* and *C. papaya* lineages, and most tested *PBS3* clade genes demonstrate functional activity, whereas genes outside this clade do not. Notably, all examined Brassicaceae species, as well as the non-Brassicaceae species *T. hassleriana* and *C. spinosa* exhibit these three traits. In contrast, *C. papaya* and all tested non-Brassicales species lack them. Although these adaptations appear to have emerged in an ancestral Brassicales species, the probability of three independent adaptations occurring simultaneously in a single species is exceedingly low. This conundrum underscores a fundamental challenge in evolutionary biology: while the evolution of a single gene is rarely sufficient to establish a new biosynthetic function, the concurrent evolution of multiple genes is statistically improbable, making the emergence of complex pathways an intriguing evolutionary puzzle. Within the IC pathway, given that *ICS* genes are broadly conserved across land plants—albeit with lower enzymatic activity in non-Brassicales species—and that promiscuous enzymatic activity could potentially catalyze the conversion of IC to SA or non-enzymatic reaction could occur (43), EDS5 neofunctionalization might be the initial key step in the evolution of IC-based SA biosynthesis. This was likely followed by the enhancement of ICS enzymatic activity and the specialized evolution of *PBS3*. Testing this hypothesis remains challenging due to the limited availability of non-Brassicaceae Brassicales genomes. Thus, expanding genomic resources within the Brassicales order will be helpful to elucidate how *ICS*, *EDS5*, and *PBS3* were recruited, adapted, and integrated into a functional biosynthetic network for IC-based SA biosynthesis.

A recent study proposed that the IC pathway evolved in the MRCA of land plants, based on the presence of homologs of *ICS*, *EDS5*, and *PBS3* in various land plant genomes (28). While the reduction of SA levels following *ICS* silencing in soybean and cotton (23, 24) supports this hypothesis, previous studies indicate that *ICS* in rice and barley does not contribute to SA biosynthesis (26, 47). Additionally, maize *ics1* mutant plants exhibit reduced PAL activity (25), suggesting that *ICS* involvement in SA accumulation may result from pleiotropic effects rather than direct participation in the IC pathway. Our analysis clearly demonstrates that only Brassicales species that diversified after the divergence of *M. oleifera* and *C. papaya* lineages exhibit enhanced ICS activity. Moreover, *EDS5* neofunctionalization for IC-based SA biosynthesis has occurred and the functional emergence of specialized *PBS3* are exclusive to those Brassicales species. Taken together, these findings strongly support the notion that the IC pathway is a derived trait that evolved within Brassicales species that diversified after the divergence of *M. oleifera* and *C. papaya* lineages.

Our structural predictions, combined with sequence alignment analyses and functional experiments, have identified key amino acid residues within the putative substrate-binding pocket of the EDS5 C-lobe that are essential for SA biosynthesis. The predicted role of EDS5 as an IC transporter, facilitating its movement from chloroplasts to the cytosol, may have been enabled by the acquisition of these residues. Through site-directed mutagenesis, we demonstrated that these amino acids are essential for EDS5 function, and notably, only functional EDS5 variants from Brassicales possess these residues. Thus, these amino acid changes in the ancestral EDS5 clade likely represent necessary adaptations for the IC pathway. Further investigation into how these amino acids influence substrate transport dynamics and the precise identity of the EDS5 substrate will be crucial for advancing our understanding of the evolution and mechanistic underpinnings of the IC pathway.

## MATERIALS AND METHODS

### Phylogenetic analysis

For the species tree used for calibration of phylogenetic tree, the CDS of various plant species were downloaded from Phytozome (https://phytozome-next.jgi.doe.gov/). *T. hassleriana*, *C. spinosa*, *A. arabicum*, *G. gynandra*, and *M. oleifera* CDS were obtained from published literature (48–52). As previously published (40), the nucleotide sequences were translated into protein sequences by EMBOSS v6.6.0.0 (https://github.com/kimrutherford/EMBOSS/), then aligned with MAFFT v7.480 (53), and trimmed by ClipKIT v0.16.0 (https://github.com/JLSteenwyk/ClipKIT). The protein maximum likelihood (ML) trees were generated using IQ-TREE v2.1.4 (https://github.com/iqtree/iqtree2) with the LG+G4 model. The collection of the CDS from various species gene trees was subjected to the coalescence-based species tree inference with ASTRAL v5.7.3 (https:// github.com/smirarab/ASTRAL). *M. polymorpha* was used as the outgroup for rooting.

For gene trees, homologs of *A. thaliana* ICS, EDS5, PBS3, and EPS1 from various species were found using tblastx v2.11.0 with parameters settings being a maximum of 5000 target sequences, cutoffs of E-value < 0.01, and > 50% query coverage, as previously published (40). The protein structure domain was predicted using rpsblast v2.11.0 with the E-value <0.01. Protein ML trees were generated with the LG+G4 model as described above. The ML trees were used as the starting gene trees for GeneRax v2.0.4 (https://github.com/BenoitMorel/GeneRax) to generate a rooted, species-tree-aware gene trees. For amino acid sequence alignment, full-length amino acid sequences were aligned using MUSCLE implemented in the Geneious 9.1.2 software at the default parameters.

### Structure prediction

The structural models of AtEDS5 and AtICS1 were predicted using the ColabFold version (54) of AlphaFold2 (55). All generated models of AtEDS5 and AtICS1 exhibited high confidence, except for their disordered N-terminal regions, as indicated by high predicted local distance difference test (pLDDT) scores and low predicted aligned error (PAE) values (Supplementary Figure S18). The pLDDT and PAE metrics reflect local accuracy and the confidence in the relative positioning of two residues within the predicted structures, respectively. For further analysis, we selected the top-ranked model from the predictions.

### Plant materials and growth conditions

*A. thaliana* Col-0, *C. rubella* N22697, and *E. salsugineum* Shandong plants were grown in a lab soil (8:4:1 peat soil: nutrient soil: vermiculite) in a chamber at 22°C/20°C under a 12/12 h (light/dark) cycle with 60% relative humidity. *N. benthamiana*), soybean (*G. max* Williams 82), and tomato plants (*S. lycopersicum* Moneymaker) were grown in the same lab soil in a chamber at 25°C under a 16/8 h (light/dark) cycle, with 60% relative humidity. Rice (*O. sativa* Kitaake) seeds were germinated on wet filter paper in a petri dish, and four- to five-day-old seedlings were transplanted to the same lab soil. Seedlings of rice and maize (*Z. mays* B73) were grown in a greenhouse at 28°C under a 16/8 h (light/dark) cycle. The *A. thaliana* accession Col-0 was the wild-type of all *A. thaliana* mutants used in this study, and *sid2-2* (56) or *eds5-1* (20) was the background of all *A. thaliana* transgenic plants.

### Genomic DNA extraction

Genome DNA was extracted from plants using DNA extraction Kit (CWBIO, CW0531M) following the manufacturer’s protocol.

### RNA extraction and cDNA synthesis

Total RNA was extracted from plants using RNA extraction Kit (Aidlab, RN3302). First-strand cDNA was synthesized from 1 µg total RNA using HiScript III RT SuperMix for qPCR with gDNA Remover (Vazyme, R323-01).

### Primers used in this study

All primers are listed in Supplementary Table S1.

### Generation of constructs for *Agrobacterium* transient assay in *N. benthamiana*

The open reading frames (without STOP codon) of *ICS*, *EDS5*, and *PBS3* homologs from *T. hassleriana* (*ThICS1a*, *ThICS1b*, *ThEDS5*, *ThEDS5H_1*, *ThEDS5H_2, ThPBS3_1*, *ThPBS3_2*, *ThPBS3_3*, and *ThPBS3_4*), *C. spinosa* (*CasICS1*, *CasEDS5*, *CasPBS3_1*, *CasPBS3_2*, *CasPBS3_3*, and *CasPBS3_4*), *C. papaya* (*CpICS1*, *CpEDS5A*, and *Cp3*), *C. sinensis* (*CisICS1*, *CisEDS5Aa*, *CisEDS5Ab*, and *Cis3*), *G. raimondii* (*GrICS1*, *GrEDS5Aa*, *GrEDS5Ab*, and *Gr3*), and *O. sativa* (*OsICS1*, and *OsEDS5A*) were synthesized by Sangon company in China. The open reading frames (without STOP codon) of *A. thaliana* (*AtICS1, AtICS2*, *AtEDS5*, *AtEDS5H*, *AtPBS3*), *C. rubella* (*CrICS1*, *CrICS2*, *CrEDS5*, *CrEDS5H*, *CrPBS3*), *E. salsugineum* (*EsICS1*, *EsICS2*, *EsEDS5H*, and *EsPBS3*), *S. lycopersicum* (*SlICS1*, *SlEDS5Aa*, *SlEDS5Ab*, and *Sl3*), *G. max* (*GmICS1a*, *GmICS1b GmEDS5Aa*, *GmEDS5Ab*, and *Gm3*), *Z. mays* (*ZmICS1*, and *ZmEDS5A*), *M. polymorpha* (*MpICS1*, and *MpEDS5A*) were PCR-amplified from cDNA using PrimeSTAR MAX DNA polymerase (Takara R045A). pENTR4 was cut using the restriction enzymes *EcoR*V *and Nco*I (TAKARA) to generate the linearized vector. The PCR fragment and the linearized pENTR4 were ligated by seamless cloning Kit (Vazyme C112-01), which were then used to transform *E. coli* DH5α that was selected with 30 µg/mL kanamycin. The sequence-verified plasmid DNA was subjected to Gateway Cloning into the destination vectors (*ICS* homologs were ligated to the pGWB617 and pGWB641 vectors, and *EDS5*, *PBS3* homologs were ligated to the pGWB641 vector) following the manufacturer’s protocol (Invitrogen, 11791-020), which were then used to transform *E. coli* DH5α that was selected with 50 µg/mL spectinomycin. *Agrobacterium tumefaciens* GV3101 was transformed with the resulting plasmid through electroporation and selected with 40 µg/mL rifampicin and 50 µg/mL spectinomycin. Amino acid point mutations in the *EDS5* and *ICS1* genes were generated through targeted mutagenesis using PCR methods as previously described (57).

### Generation of *A. thaliana* transgenic plants

The *AtICS1* and *AtEDS5* promoters were amplified from Col-0 genome DNA using PrimeSTAR MAX DNA polymerase (Takara R045A). pENTR4 was cut using the restriction enzymes *EcoR*V and *Nco*I (TAKARA) to generate the linearized vector. The PCR fragment and the linearized to pENTR4 were ligated by seamless cloning Kit (Vazyme C112-01), which were then used to transform *E. coli* DH5α that was selected with 30 μg/mL kanamycin. The plasmid DNA was extracted and cut with *EcoR*V and *Nco*I (TAKARA) to linearize the vector. The coding sequences (with STOP codon) were amplified from the cDNA of the respective plants or synthesized as described above. The PCR fragment and the linearized pENTR4 containing the promoter were ligated by seamless cloning Kit (Vazyme C112-01), which were then used to transform *E. coli* DH5α that was selected with 30 μg/mL kanamycin. The sequence-verified plasmid DNA was subjected to Gateway Cloning (Invitrogen, 11791-020) into pGWB501, which was then used to transform *E. coli* DH5α that was selected with 50 µg/mL spectinomycin. *A. tumefaciens* GV3101 was transformed with the resulting plasmid through electroporation and selected with 40 µg/mL rifampicin and 50 µg/mL spectinomycin. The transformed *A. tumefaciens* was used to transform *A. thaliana sid2-2* or *eds5-1* plants by floral dipping (58). T1 plants were selected on half-strength MS medium containing 20 µg/mL hygromycin and 50 µg/mL cefotaxime. T2 or T3 transgenic plants were used for the subsequent experiments.

### Selection of maize *eds5* mutant

The maize *eds5A* mutant seeds, generated with ethyl methanesulfonate (EMS) mutagenesis, were obtained from the website www.elabcaas.cn. After sowing seeds and growing the seedlings, DNA was extracted, and *ZmEDS5A* gene was amplified, followed by Sanger sequencing to screen desired homozygotes for subsequent experiments.

### Generation of *eds5A* mutant of tomato, soybean, and rice with CRISPR/Cas9

For generating tomato *eds5A* mutant, CRISPR/Cas9 binary vectors (pTX) was used, which contains a tomato U6 promoter and 35S promoter to drive gRNA and Cas9 (59), respectively. The gRNA that targets *SlEDS5A* was cloned into the BsaI site of pTX vector. The tomato moneymaker was used for transformation as previously described (60). T0 generation explants were further screened on the medium containing kanamycin.

For generating soybean *eds5A* mutants, CRISPR/Cas9 construct was generated as previously described (61). Briefly, target site-containing primers were annealed and cloned into the sgRNA expression cassettes pYLsgRNA-AtU3b, pYLsgRNA-AtU3d, pYLsgRNA-AtU6-1, and pYLsgRNA-AtU6-29 at a BsaI site. Nested PCR was employed to amplify the integrated sgRNA expression cassettes, which were then ligated into the pYSLCRISPR/Cas9 vector to obtain the GmEDS5Aa/b-Cas9 construct. Subsequently, *A. tumefaciens* strain EHA101 was transformed with GmEDS5Aa/b-Cas9 and used for transformation of Williams 82 plants via cotyledon-node method as previously described (62). Extract genomic DNA from the regenerated plantlets and sequence the PCR products to confirm mutations. Transfer the confirmed mutant plants to soil for further analysis of segregation patterns in subsequent generations. Homozygous plants were used for further experiments.

For generating rice *eds5A* mutants, four target sites in the *OsEDS5A* region were designed using CRISPR-P2.0 software. The genome editing plasmids were constructed as previously described(63). Briefly, for each gRNA, a pair of DNA oligos with 20 bp targeting sequences and appropriate 4 nt overhangs were synthesized by Sangon company in China and then annealed to form a dsDNA duplex. Then resulting dsDNA oligos were subsequently inserted into the BsaI sites of pRGEB32B1, B4, B7, and B10 plasmids, respectively. These Cas9/gRNA plasmid constructs were then transformed into rice Kitaake calli by ToWin Biotechnology company in China. T0 generation explants were screened on media containing hygromycin, and DNA was extracted for PCR to verify mutations. Seedlings with confirmed mutations were transferred to soil for continued growth. Once mature, the seeds were reintroduced to the soil for further screening, and homozygous plants were used for subsequent experiments.

### Agrobacterium transient assay in N. benthamiana

*A. tumefaciens* GV3101 strains harboring different constructs were grown in YEB medium containing 40 µg/mL rifampicin and 50 µg/mL spectinomycin, and then resuspended in infiltration buffer (10 mM MES (pH 5.6), 10 mM MgCl_2_, and 150 µM acetosyringone) to an OD_600_ of 0.3. These *A. tumefaciens* strains were incubated in the dark at room temperature on a rotary shaker set to 150 rpm for 2-2.5 h. In some cases, multiple *A. tumefaciens* strains were mixed in equal concentrations to achieve a final OD_600_ of 0.3. All *A. tumefaciens* suspension for infiltration included *A. tumefaciens* carrying P19 at the final ratio of 10%. The resulting suspension was then infiltrated into the leaves of 4 to 6-week-old *N. benthamiana* plants using a needleless syringe.

### Salicylic acid measurements

Infiltrated *N. benthamiana* leaves were harvested and rapidly frozen in liquid nitrogen at 5 days post-infiltration (dpi). *Arabidopsis* transgenic lines were infiltrated with *P. syringae* DC3000 (OD_600_=0.002), the samples were collected at 1 dpi and the weight of each sample was recorded. Soybean plants were infiltrated with *P. syringae Psg*neau001 (OD_600_=0.5), the samples were collected at 2 dpi and the weight of each sample was record. 3-week-old maize plants, and 4-week-old rice plants were collected, and the weight of each sample was recorded. The samples (approximately 100 mg) were thoroughly homogenized in the TissueLyser at 60 Hz for 1 minute. After this, 500 µL of 50% methanol containing an internal standard (SA-d4) was introduced into each tube for normalization. The samples were centrifugated at 4°C and 13000 ×g for 5 minutes, followed by filtration of the supernatant using a 0.22 μm filter. The filtered extracts were then analyzed using a TSQ Quantiva and TSQ Endura Altis Quantis (Thermo Fisher). The column (Waters) temperature was maintained at 40°C with a 0.4 ml min^-1^ flow rate, employing a mobile phase gradient of water + 0.1% formic acid (A) and acetonitrile + 0.1% formic acid (D) as follows: 0-0.5 min 2% D; 0.5-6 min 98% D; 6-8 min 98% D; 8-8.1 min 2% D; 8.1-10 min 2% D. During the analysis, the following MRM transitions were monitored: SA (m/z 137→93), SA-d4 (m/z 141→97), and SAG (m/z 299→137). Peak selection and integration of acquired MRM data files were done using FreeStyle^TM^ 1.3 SP2 software (Thermo Scientific).

### Subcellular localization

Leaf sections from the infiltrated *N. benthamiana* plants at 2 dpi were delicately excised and placed onto glass slides. YFP was excited using a 488 nm wavelength and detected at 514 nm, while chlorophyll auto-fluorescence was detected at 650-750 nm using confocal laser scanning microscopy LEICA SP8. Merging of images was performed using LAS X software.

### qRT-PCR

Total RNA was extracted followed by cDNA synthesis as described above, qPCR was performed on a Real-Time PCR system (Biorad). Cq values were calculated by subtracting the Cq value of the target gene from that of the reference gene. These ΔCq values were considered to be log_2_ expression values. The reference gene: *AtActin* (*A. thaliana*), *CrActin* (*C. rubella*), *EsActin* (*E. salsugineum*), *GmELF1b* (soybean), *SlActin* (tomato) *ZmUBQ* (maize), *OsActin* (rice).

### Immunoblotting

Ground plant tissues were thoroughly mixed with extraction buffer consisting of 50 mM Tris-HCl (pH=7.5), 150 mM NaCl, 10 mM MgCl_2_, 1 mM EDTA, 10 mM NaF, 2 mM Na_3_VO_4_, 25 mM β-sodium glycerophosphate, 10% glycerol, and 0.1% NP40. The mixture was incubated on ice for 10 min and then heated at 70°C for 5 min (no heating for EDS5). After centrifugation to remove debris, proteins were separated by SDS-PAGE and transferred to a PVDF membrane using a semi-dry transfer system (Biorad). The transferred blot was incubated in 1×TBST supplemented with 5% skim milk for 1 h, followed by incubation with anti-Myc or anti-GFP antibodies at 4°C overnight. Signals were visualized using a chemiluminescent substrate and protein bands were detected using an imaging system (Tanon).

### Statistical analysis

All quantitative values were log-transformed to meet the criteria for statistical analyses using Student’s t-test.

## ACKNOWLEDGMENTS

This work was supported by the National Natural Science Foundation of China (32170298 and 32250710139 to K.T.), the National Key R&D Program of China (2022YFA1304400 to K.T. and X.H.), and the United States Department of Agriculture-National Institute of Food and Agriculture (grant no. 2020-67013-31187 to F.K.). We thank Masahito Nakano for providing technical assistance, Xiaoxia Wu for providing *Pseudomonas syringae* Nequ001, and Guotian Li for providing the *Xanthomonas oryza* PXO99.

## AUTHOR CONTRIBUTIONS

K.H. and K.T. designed research; K.H., Y.T., L.J., W.K., Y.W., L.Z., P.L., J.H., W.J., R.H., H.M., F.K., and M.H. performed research; H.N., B.L., X.L., K.X., K.F., and L.G. contributed new reagents/ analytic tools; K.H., F.K., X.H., and K.T analyzed data; and K.H., F.K., and K.T. wrote the paper.

---

**Figure S1.**
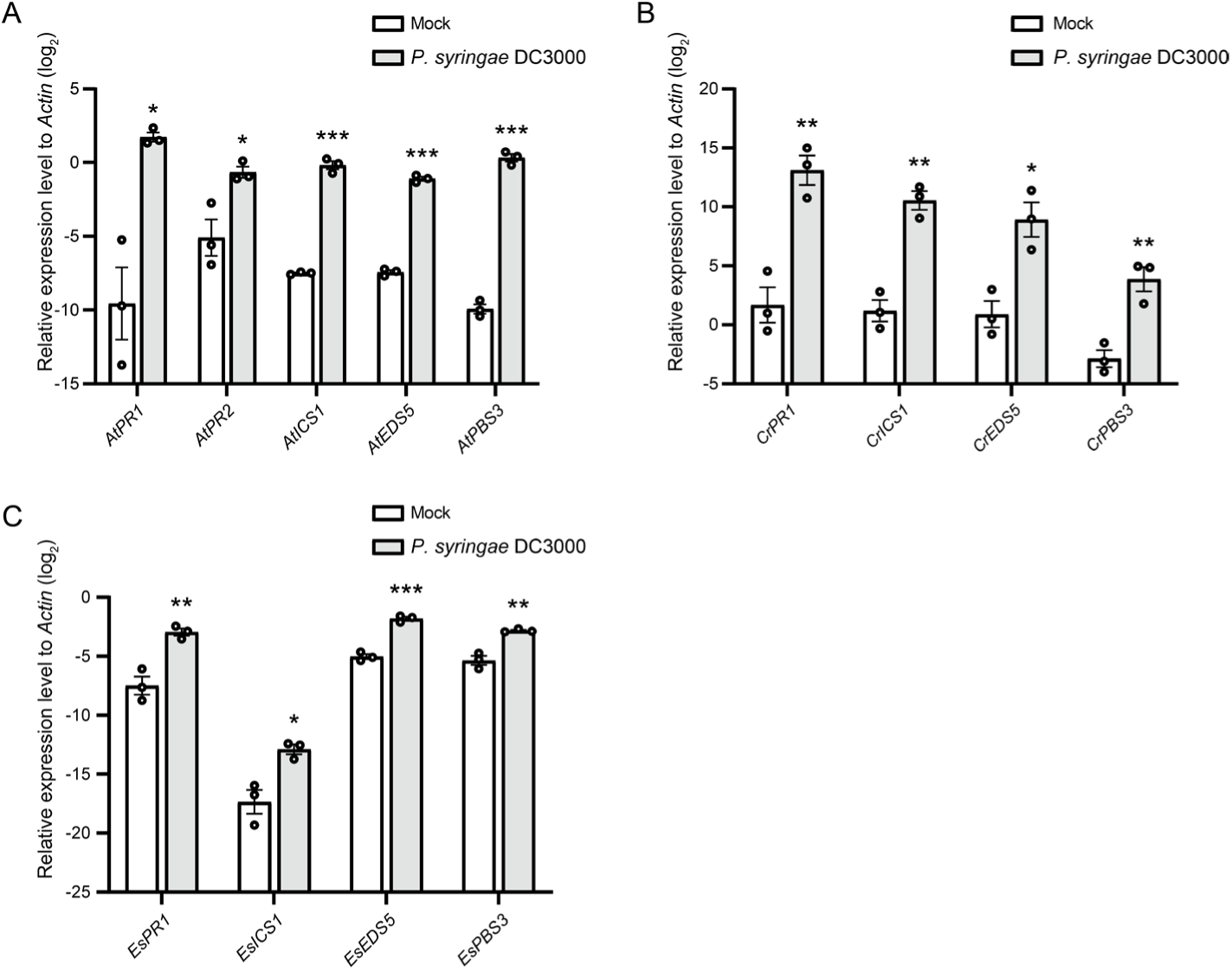
*ICS1*, *EDS5*, and *PBS3* expression is induced following pathogen infection in Brassicaceae plants. (**A**) Expression levels of *AtPR1*, *AtPR2*, *AtICS1*, *AtEDS5*, and *AtPBS3* in 33-day-old *A. thaliana* (Col-0) leaves at 24 h post-infiltration with *P. syringae* DC3000 (OD_600_ = 0.01) or mock, as determined by qRT-PCR. (**B**) Expression levels of *CrPR1*, *CrICS1*, *CrEDS5*, and *CrPBS3* in 6-week-old *C. rubella* (N22697) leaves at 24 h post-infiltration with *P. syringae* DC3000 (OD_600_ = 0.01) or mock, as determined by qRT-PCR. (**C**) Expression levels of *EsPR1*, *EsICS1*, *EsEDS5*, and *EsPBS3* in 6-week-old *E. salsugineum* (Shandong) leaves at 24 h post-infiltration with *P. syringae* DC3000 (OD_600_ = 0.01) or mock, as determined by qRT-PCR. (**A-C**) Values represent relative log_2_ expression levels normalized to *Actin*. Data represent means ± SEM from 3 biological replicates. Asterisks indicate significant differences compared to mock (\**P <* 0.05, \*\**P <* 0.01, and \*\*\**P* < 0.001, two-tailed Student’s t-tests).

**Figure S2.**
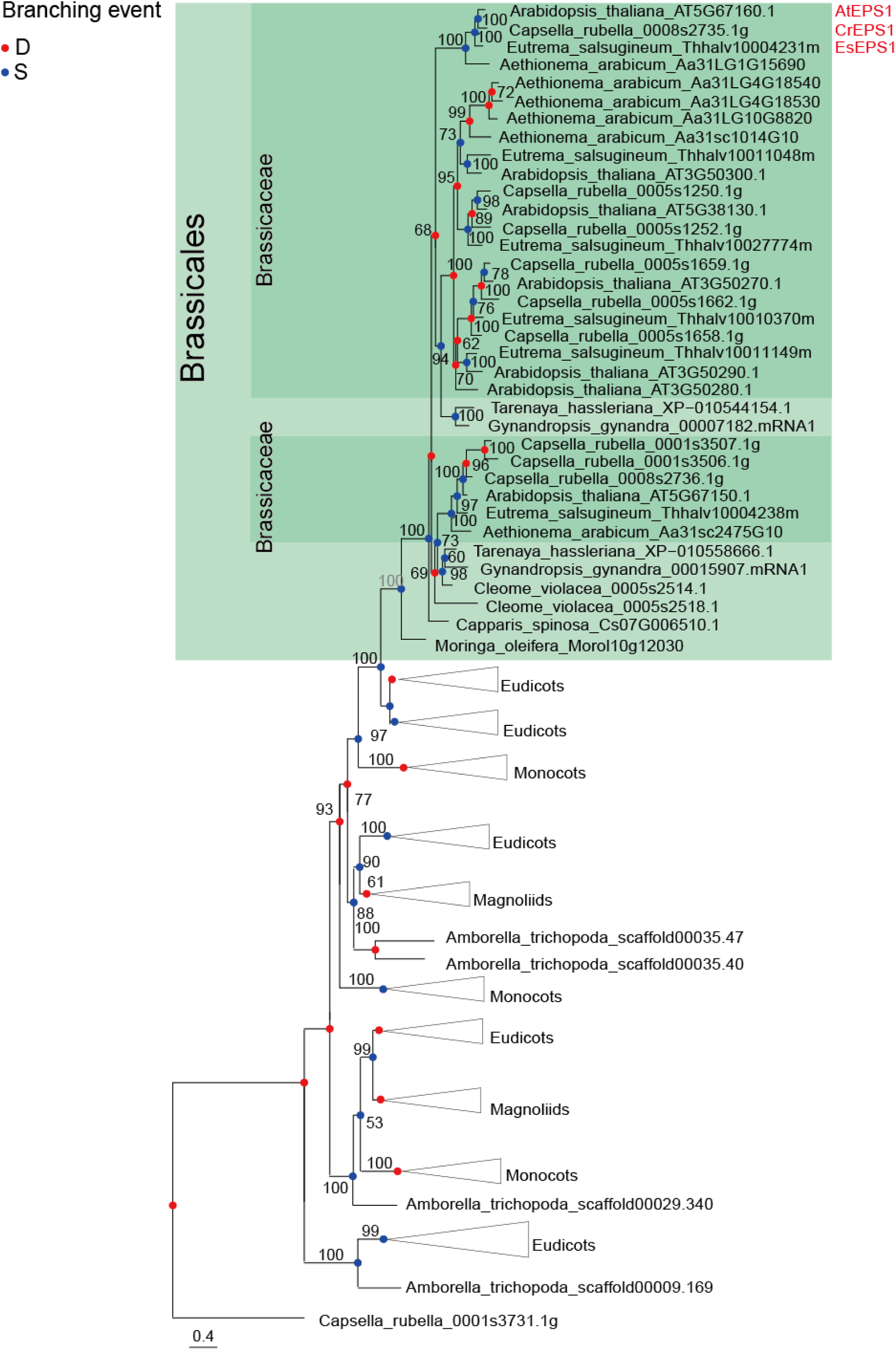
Phylogenic analysis of EPS1 homologs across plant species. The phylogenic tree of EPS1 homologs was constructed based on full-length amino acid sequences using the maximum likelihood method. Triangles represent collapsed branches. The scale bar denotes a phylogenetic distance of 0.4 amino acid substitutions per site. Bootstrap values, derived from 1000 replications, are indicated at branch nodes. “D” represents Duplication, “S” represents Speciation.

**Figure S3.**
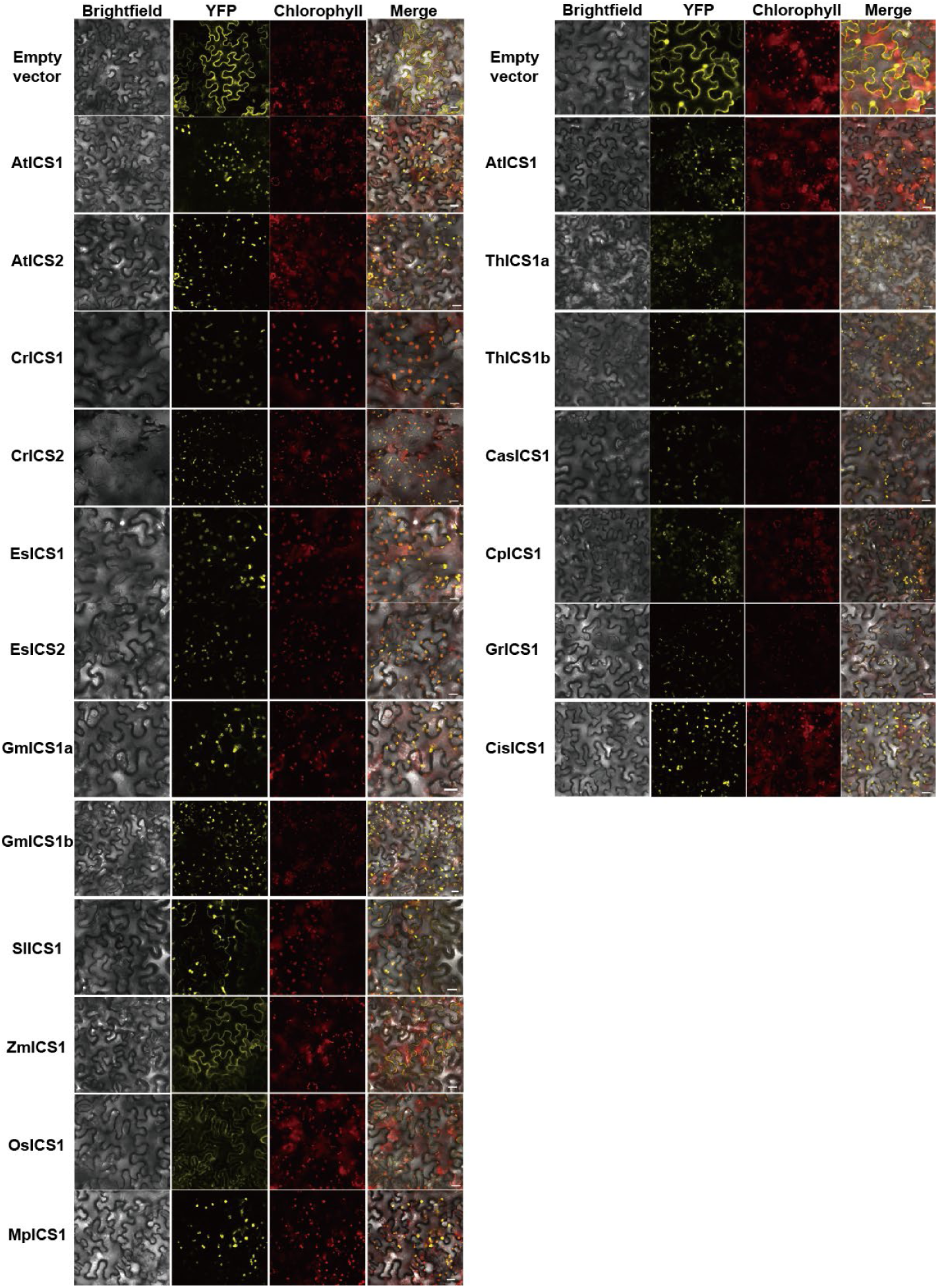
Subcellular localization of ICS from various species in *N. benthamiana*. *Agrobacterium* strains carrying *ICS-YFP* were infiltrated into *N. benthamiana* leaves. ICS subcellular localization in epidermal cells expressing ICS-YFP was analyzed by confocal microscopy at 2 dpi. Scale bars, 20 µm.

**Figure S4.**
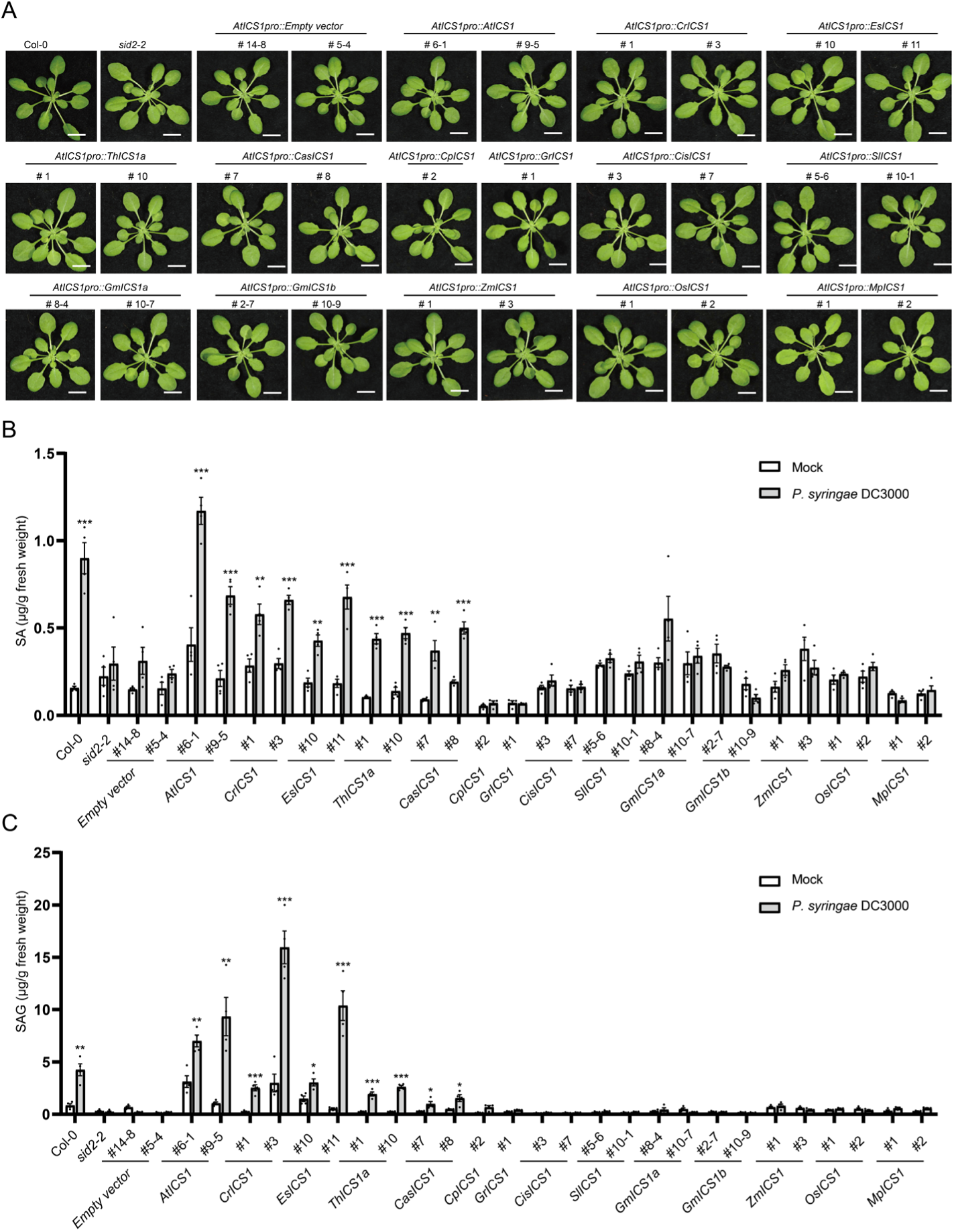
*ICSs* from only Brassicales species, except for *C. papaya*, complement the *A. thaliana sid2* mutant. (**A**) Phenotypes of 5-week-old *A. thaliana* wild type (Col-0) and transgenic *sid2* mutant plants expressing *ICS* homologs from various plant species under the control of *A. thaliana ICS1* promoter. (**B** and **C**) Leaves of 5-week-old Col-0 and transgenic plants were infected with *P. syringae* DC3000 (OD_600_=0.002) or mock. SA and SAG levels in leaf extracts were quantified at 1 dpi. Data represent means ± SEM from 4 biological replicates. Asterisks indicate significant differences compared to mock (\**P <* 0.05, \*\**P <* 0.01, and \*\*\**P* < 0.001, two-tailed Student’s t-tests).

**Figure S5.**
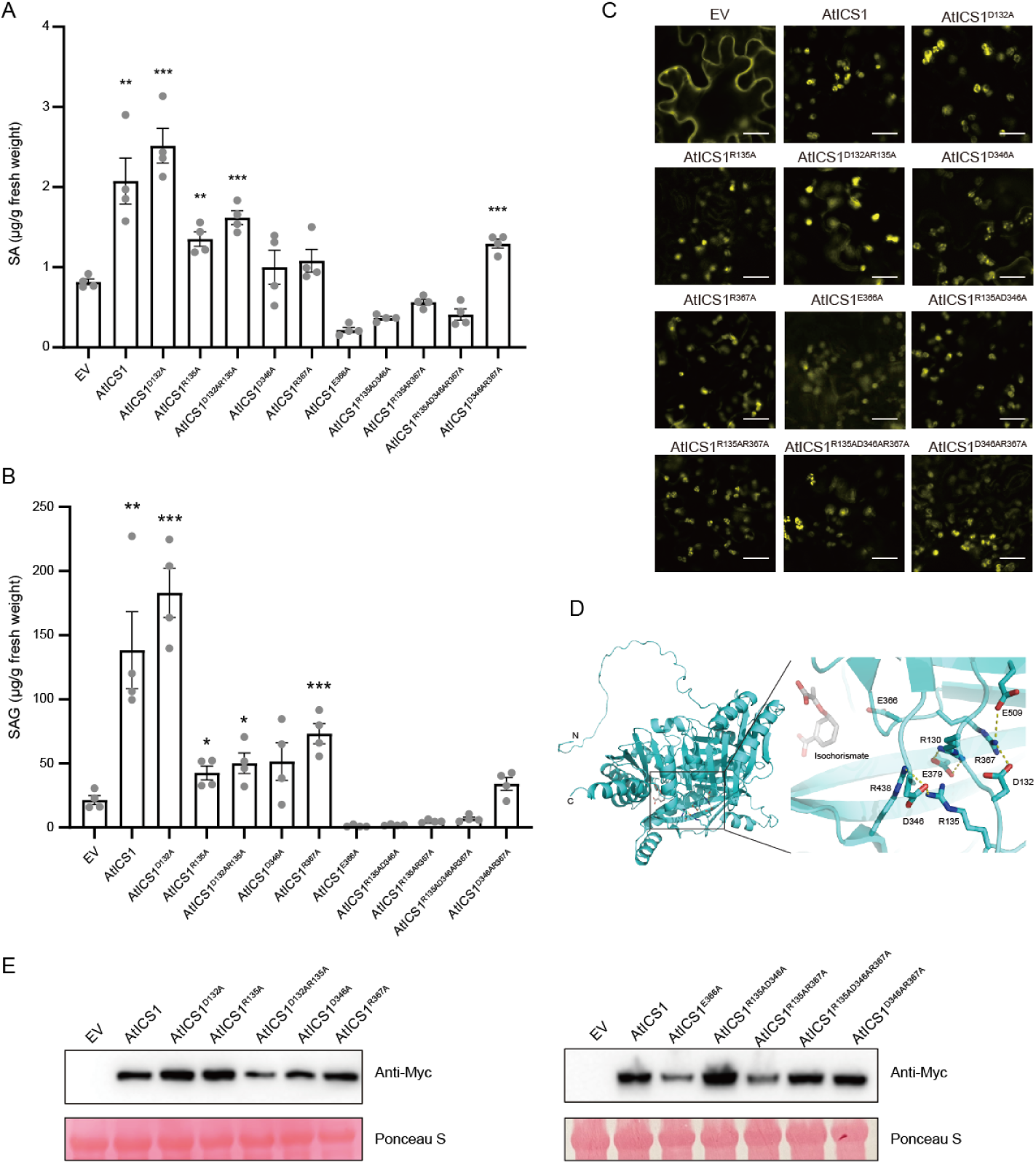
Characterization of *A. thaliana* ICS1 mutants. (**A** and **B**) *Agrobacterium* strains carrying *A. thaliana ICS-Myc* (or mutated *ICS-Myc*), *EDS5-YFP*, and *PBS3-YFP* were co-infiltrated into leaves of 4 to 6-week-old *N. benthamiana* plants. SA and SAG levels in leaf extracts were quantified at 5 dpi. Data represent means ± SEM from 4 biological replicates. Asterisks indicate significant differences compared to EV (\**P <* 0.05, \*\**P <* 0.01, and \*\*\**P* < 0.001, two-tailed Student’s t-tests). (**C**) *Agrobacterium* strains carrying *ICS1-YFP* were infiltrated into *N. benthamiana* leaves. ICS1 subcellular localization in epidermal cells was analyzed by confocal microscopy at 2 dpi. Scale bars, 20 µm. (**D**) Structural model of *A. thaliana* ICS1 predicted by AlphaFold2. Both the overall structure and a close-up view of the salt bridge network are shown. Putative hydrogen bonds are represented by dotted lines. IC from the EntC structure (PDB ID: 3HWO) was superimposed onto the predicted AtICS1 structure. (**E**) *Agrobacterium* strains carrying *ICS-YFP* were infiltrated into *N. benthamiana* leaves. Total protein was extracted from the infiltrated leaves at 2 dpi, and ICS protein expression was assessed by immunoblotting using an anti-Myc antibody.

**Figure S6.**
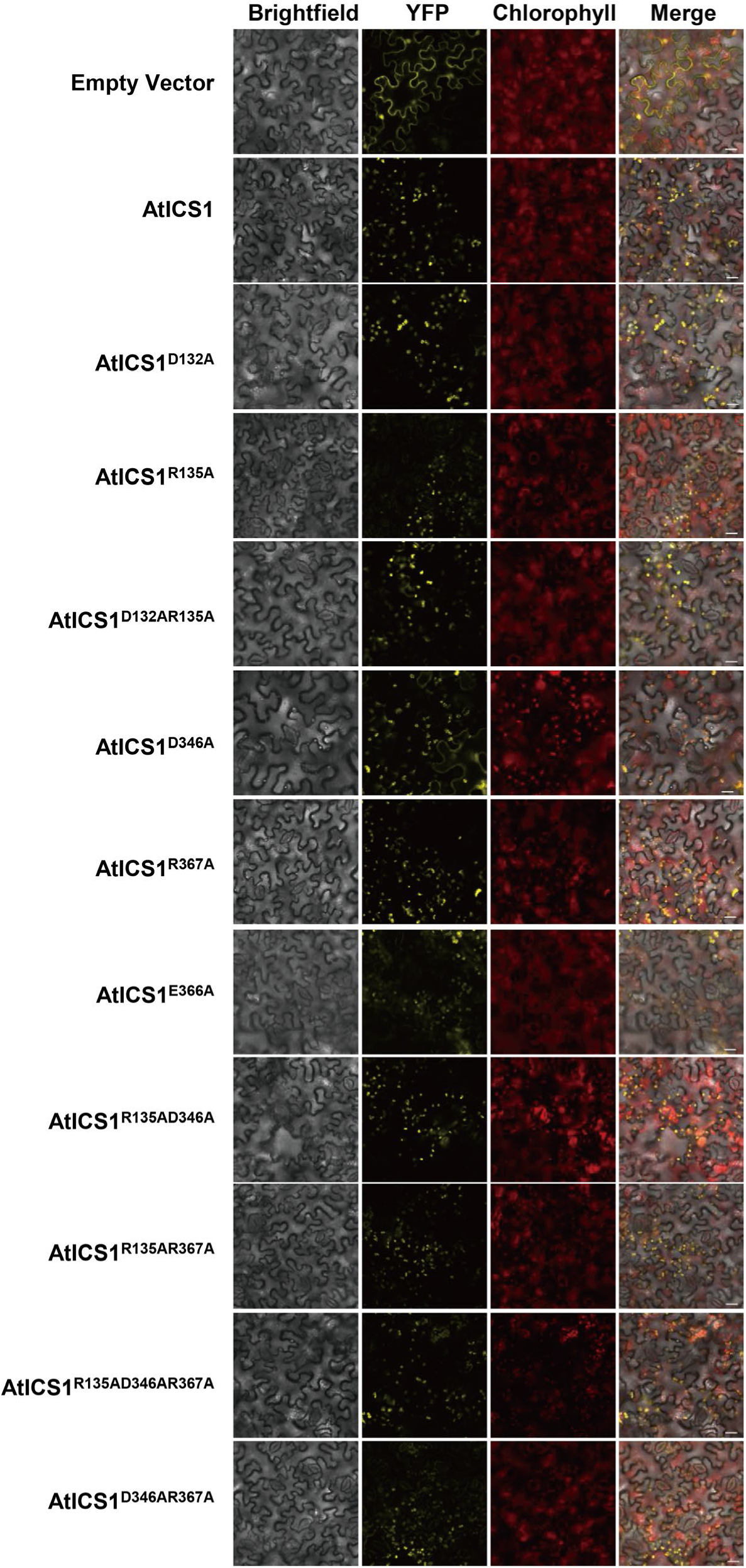
Subcellular localization of AtICS1 mutants in *N. benthamiana*. *Agrobacterium* strains carrying *AtICS1 mutants-YFP* were infiltrated into *N. benthamiana* leaves. Subcellular localization of AtICS1 mutants in epidermal cells expressing AtICS1 mutants-YFP was analyzed by confocal microscopy at 2 dpi. Scale bars, 20 µm.

**Figure S7.**
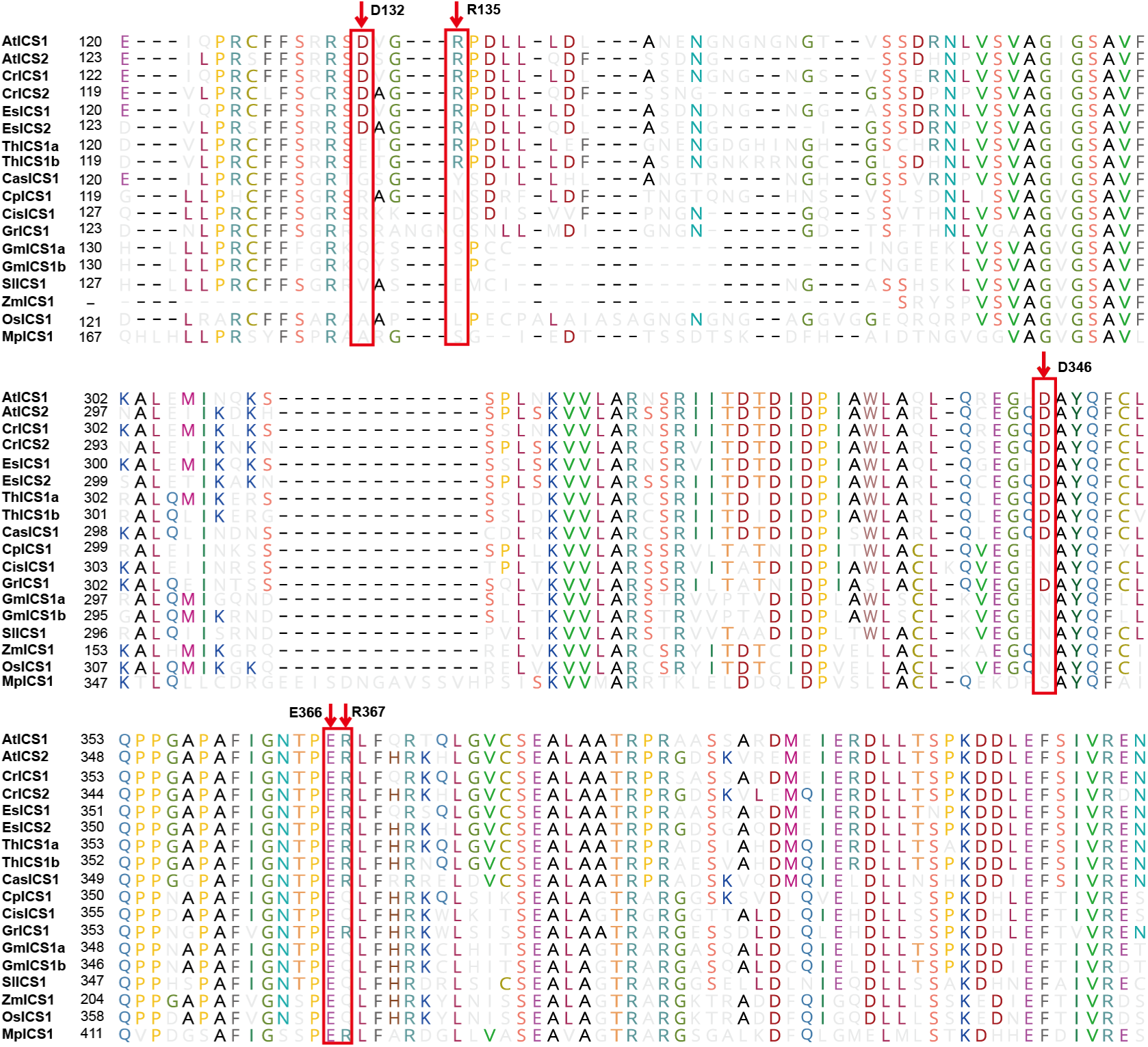
Alignment of ICS homolog amino acid sequences from various plant species. Multiple sequence alignment of ICS homologs from various plant species. Red arrows indicate the amino acid residues in *A. thaliana* ICS1 that were mutated in this study.

**Figure S8.**
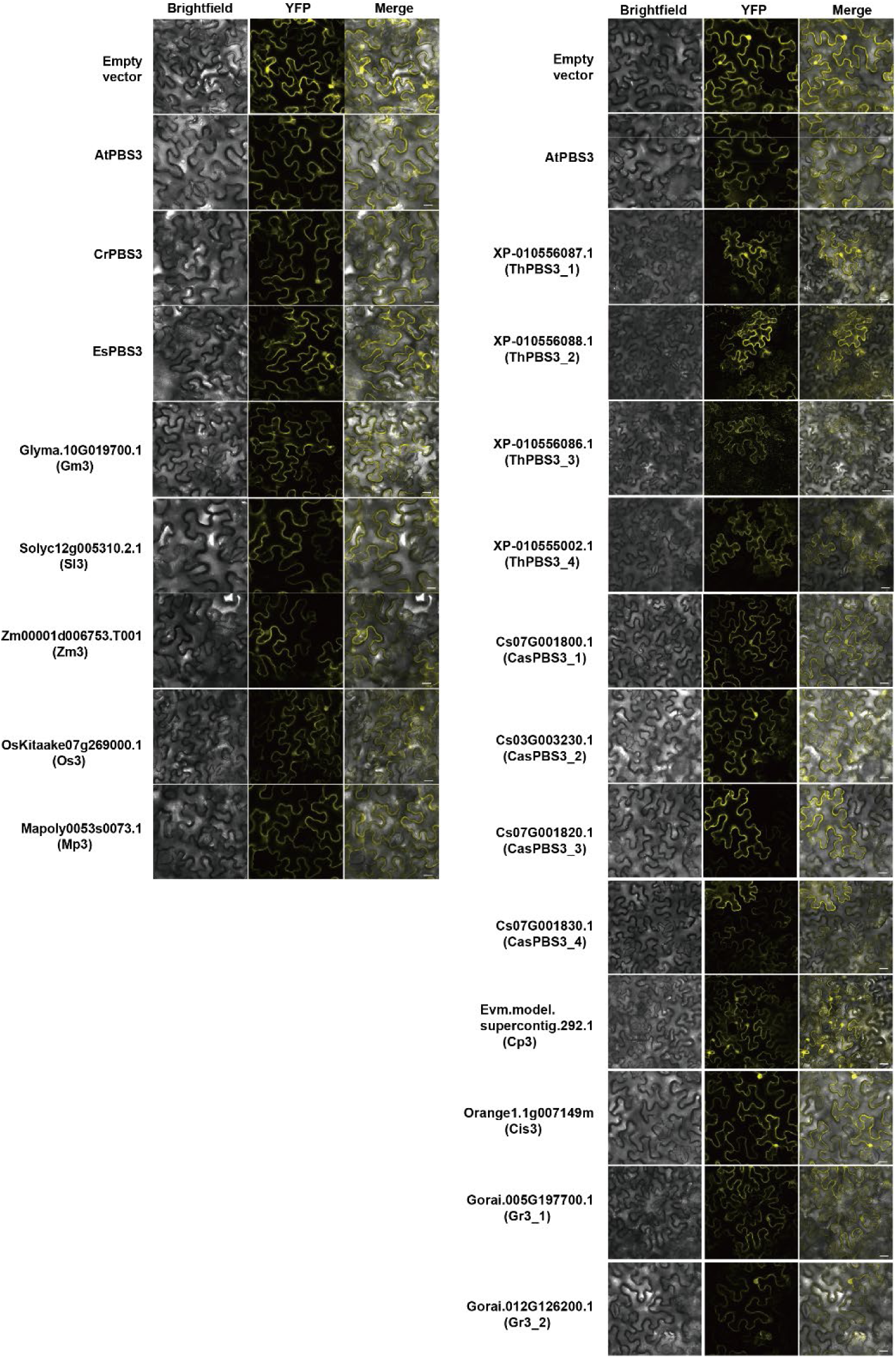
Subcellular localization of PBS3 homologs from various species in *N. benthamiana*. *Agrobacterium* strains carrying *PBS3-YFP* were infiltrated into *N. benthamiana* leaves. PBS3 subcellular localization in epidermal cells expressing PBS3-YFP was analyzed by confocal microscopy at 2 dpi. Scale bars, 20 µm.

**Figure S9.**
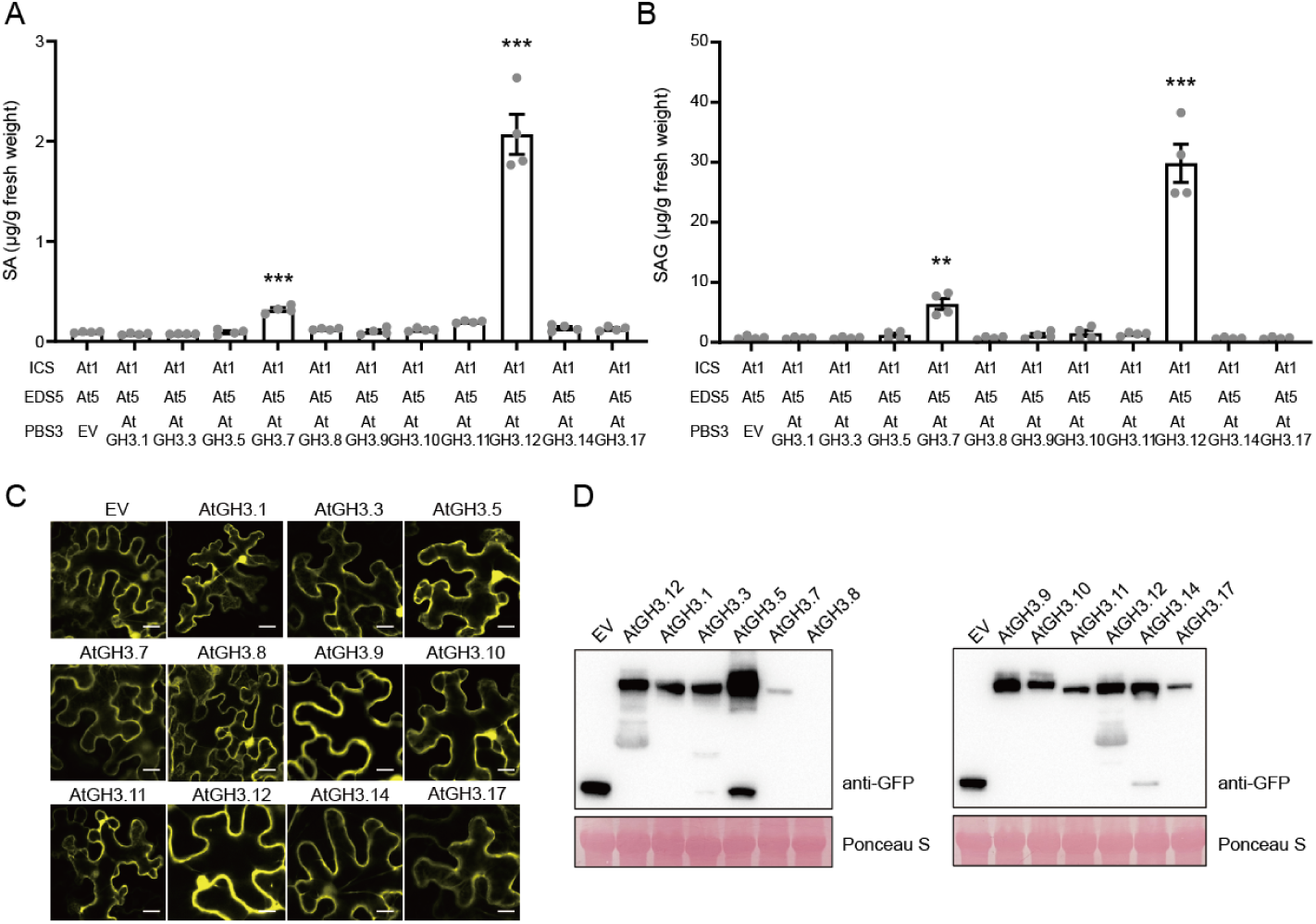
AtPBS3 (GH3.12) and GH3.7, but not other GH3 family members are functional for SA biosynthesis in *N. benthamiana*. *Agrobacterium* strains carrying *A. thaliana GH3-YFP* were co-infiltrated with *Agrobacterium* strains carrying *A. thaliana ICS-Myc* and *EDS5-YFP* into leaves of 4 to 6-week-old *N. benthamiana* plants. (**A** and **B**) SA and SAG levels in leaf extracts were quantified at 5 dpi. EV represents the empty vector control. Data represent means ± SEM from 4 biological replicates. Asterisks indicate significant differences compared to EV (\**P <* 0.05, \*\**P <* 0.01, and \*\*\**P* < 0.001, two-tailed Student’s t-tests). (**C**) *Agrobacterium* strains carrying GH3-YFP were infiltrated into *N. benthamiana* leaves. GH3 subcellular localization in epidermal cells expressing GH3-YFP was analyzed by confocal microscopy at 2 dpi. Scale bars, 20 µm. (**D**) *Agrobacterium* strains carrying *GH3-YFP* were infiltrated into *N. benthamiana* leaves. Total protein was extracted from the infiltrated leaves at 2 dpi, and GH3 protein expression was assessed by immunoblotting using an anti-GFP antibody.

**Figure S10.**
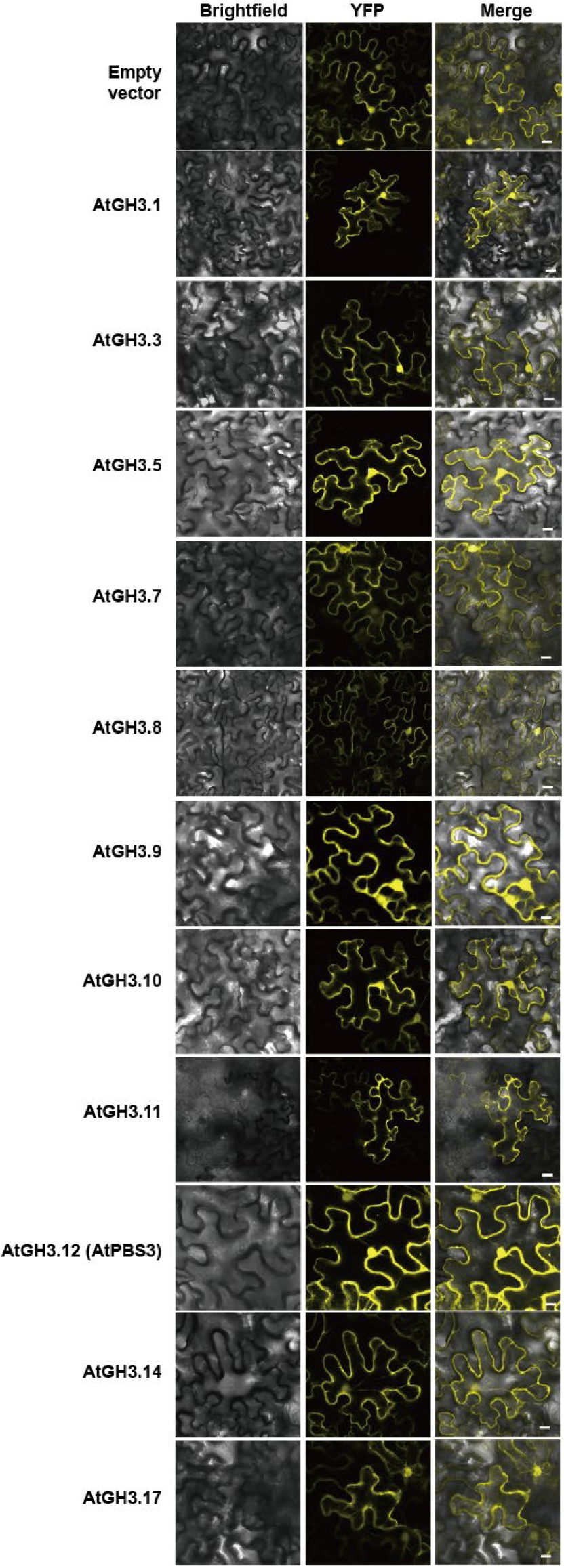
Subcellular localization of *A. thaliana* GH3s in *N. benthamiana*. *Agrobacterium* strains carrying *GH3-YFP* were infiltrated into *N. benthamiana* leaves. GH3 subcellular localization in epidermal cells expressing GH3-YFP was analyzed by confocal microscopy at 2 dpi. Scale bars, 20 µm.

**Figure S11.**
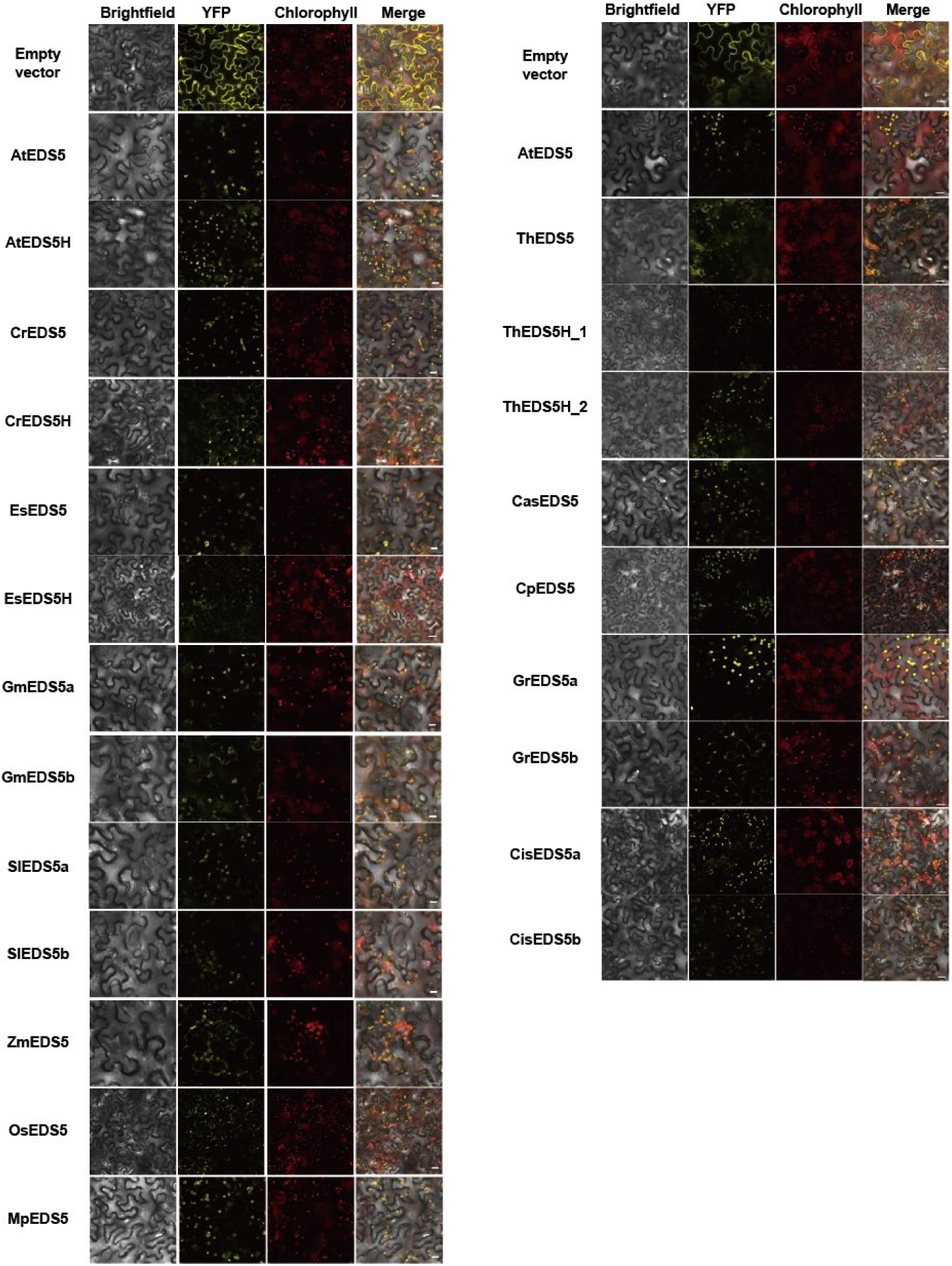
Subcellular localization of EDS5 homologs from various species in *N. benthamiana*. *Agrobacterium* strains carrying *EDS5-YFP* were infiltrated into *N. benthamiana* leaves. EDS5 subcellular localization in epidermal cells expressing EDS5-YFP was analyzed by confocal microscopy at 2 dpi. Scale bars, 20 µm.

**Figure S12.**
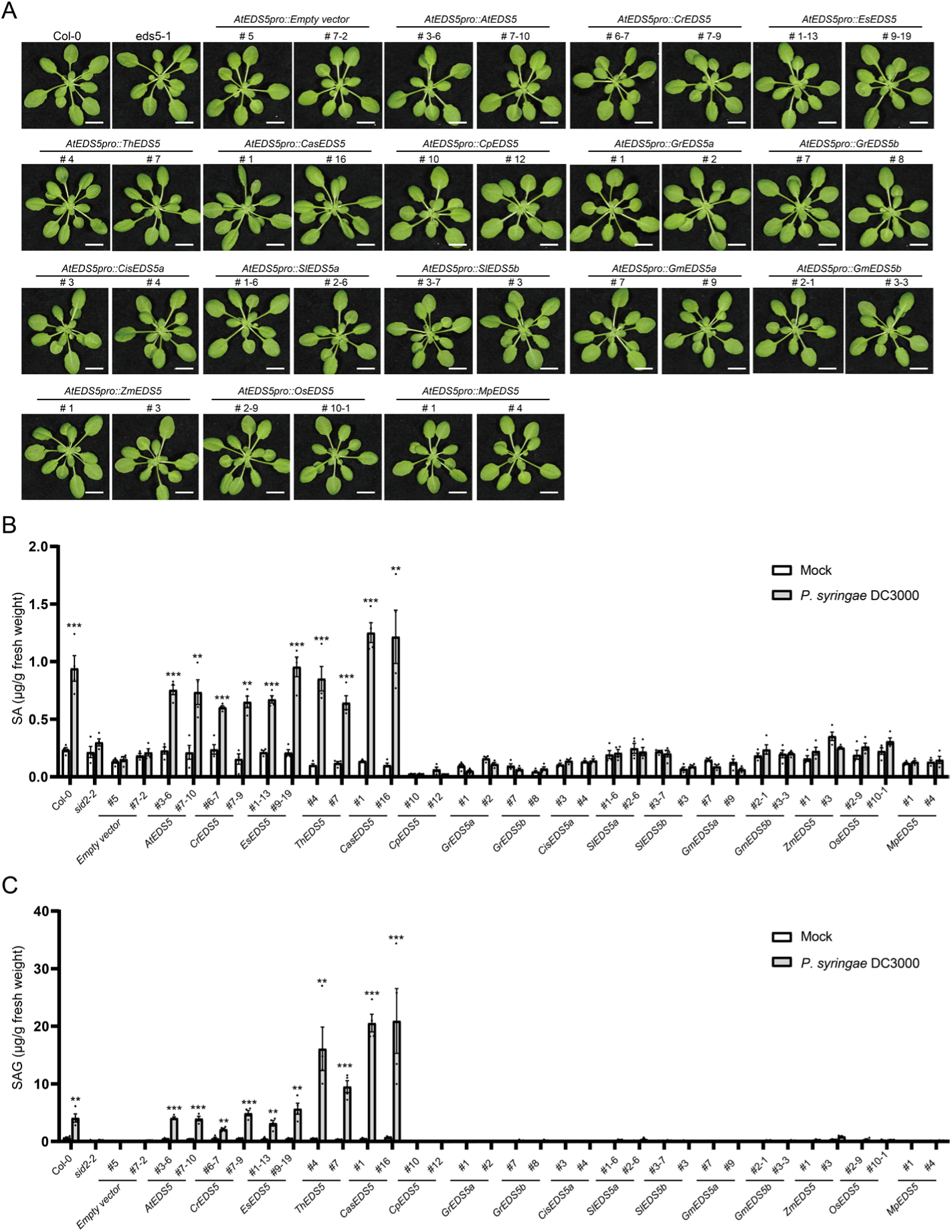
*EDS5* homologs from only Brassicales species, except for *C. papaya*, complement the *A. thaliana eds5* mutant. (**A**) The phenotype of 5-week-old *A. thaliana* wild type (Col-0), and the transgenic plants carrying homologs of *EDS5* from various plant species driven by *AtEDS5* promoter in the *eds5* mutant background. (**B** and **C**) Leaves of five-week-old Col-0 and transgenic plants were infected with *P. syringae* DC3000 (OD_600_ = 0.002) or mock. SA and SAG levels in leaf extracts were quantified at 1 dpi. Data represent means ± SEM from 4 biological replicates. Asterisks indicate significant differences compared to mock (\**P <* 0.05, \*\**P <* 0.01, and \*\*\**P* < 0.001, two-tailed Student’s t-tests).

**Figure S13.**
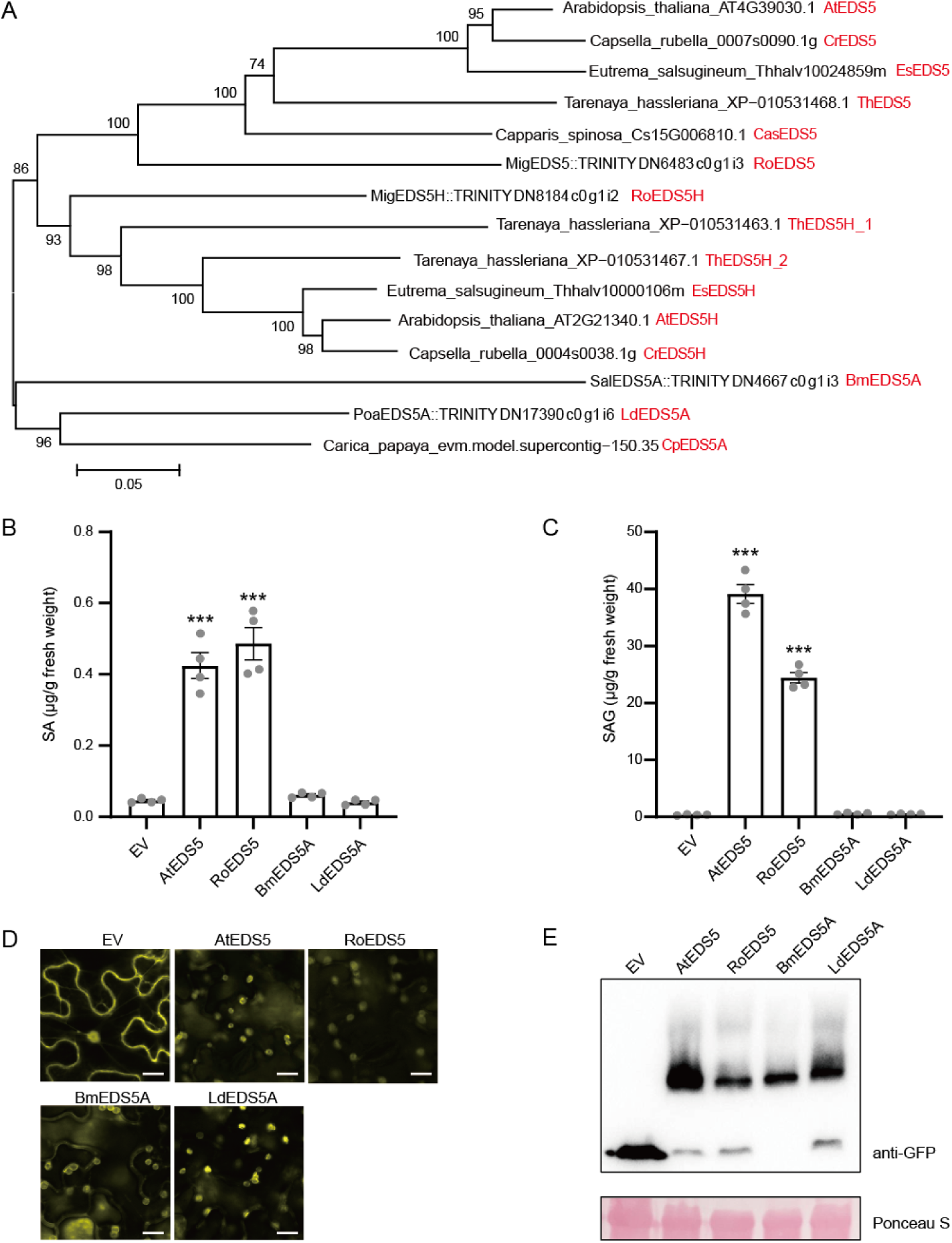
EDS5 from the Brassicales *Reseda odorata* functions for SA biosynthesis in *N. benthamiana*. (**A**) Phylogenetic trees of *EDS5* homologs from various plant species. Trees were generated based on full-length coding sequences using neighbor-joining method. Bootstrap values, derived from 1000 replications, are indicated at branch nodes. *Agrobacterium* strains carrying *EDS5-YFP* from Brassicales species were co-infiltrated with *Agrobacterium* strains carrying *A. thaliana ICS-Myc* and *PBS3-YFP* into leaves of 4 to 6-week-old *N. benthamiana* plants. (**B and C**) SA and SAG levels in leaf extracts were quantified at 5 dpi. EV represents the empty vector control. Data represent means ± SEM from 4 biological replicates. Asterisks indicate significant differences compared to EV (\**P <* 0.05, \*\**P <* 0.01, and \*\*\**P* < 0.001, two-tailed Student’s t-tests). (**D**) *Agrobacterium* strains carrying *EDS5-YFP* were infiltrated into *N. benthamiana* leaves. EDS5 subcellular localization in epidermal cells expressing *EDS5-YFP* was analyzed by confocal microscopy at 2 dpi. Scale bars, 20 µm (**E**) *Agrobacterium* strains carrying *EDS5-YFP* were infiltrated into *N. benthamiana* leaves. Total protein was extracted from the infiltrated leaves at 2 dpi and EDS5 protein expression was assessed by immunoblotting using an anti-GFP antibody.

**Figure S14.**
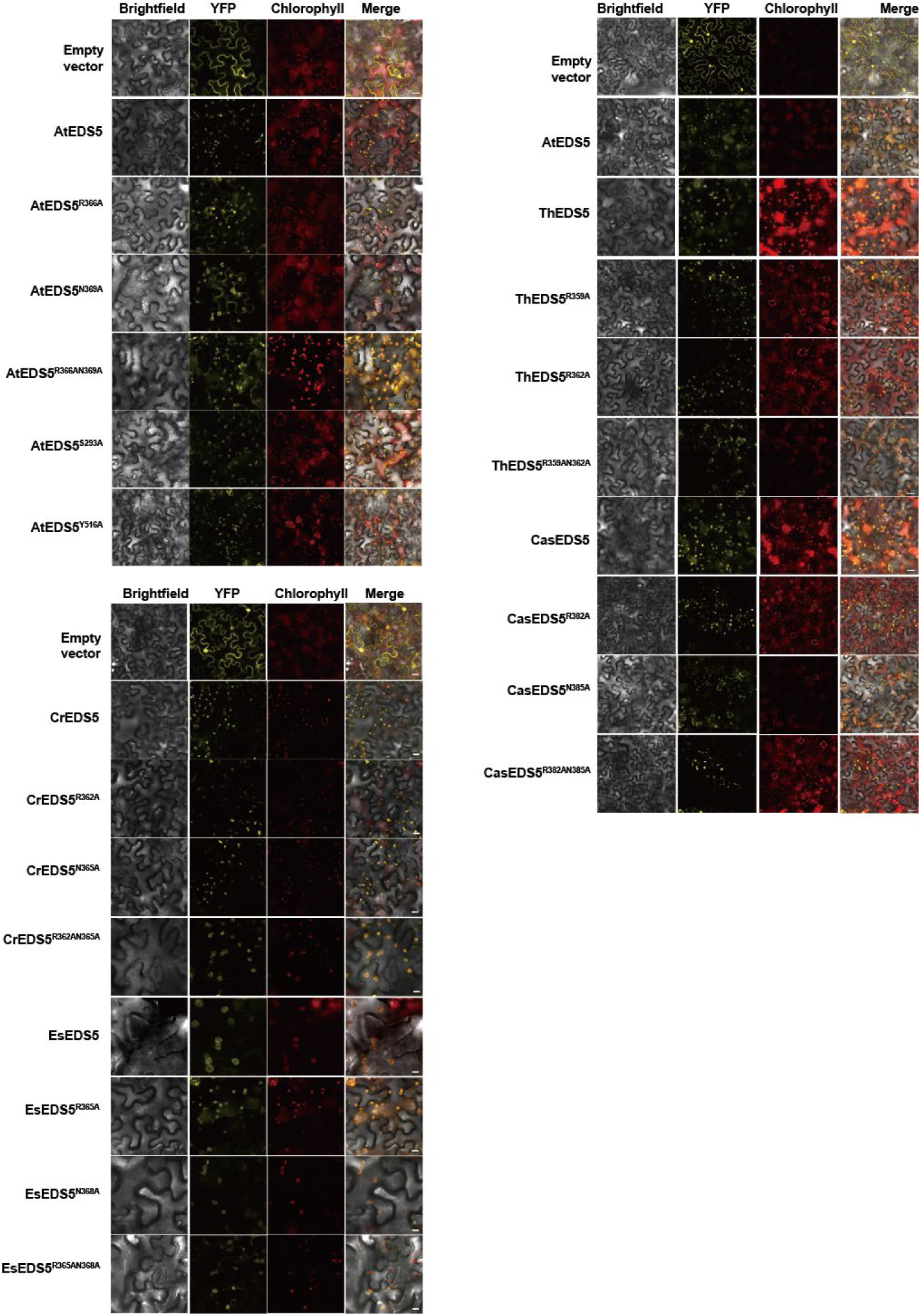
Subcellular localization of EDS5 mutants in *N. benthamiana*. *Agrobacterium* strains carrying *EDS5 mutants-YFP* were infiltrated into *N. benthamiana* leaves. EDS5 mutants subcellular localization in epidermal cells expressing AtEDS5 mutants-YFP was analyzed by confocal microscopy at 2 dpi. Scale bars, 20 µm.

**Figure S15.**
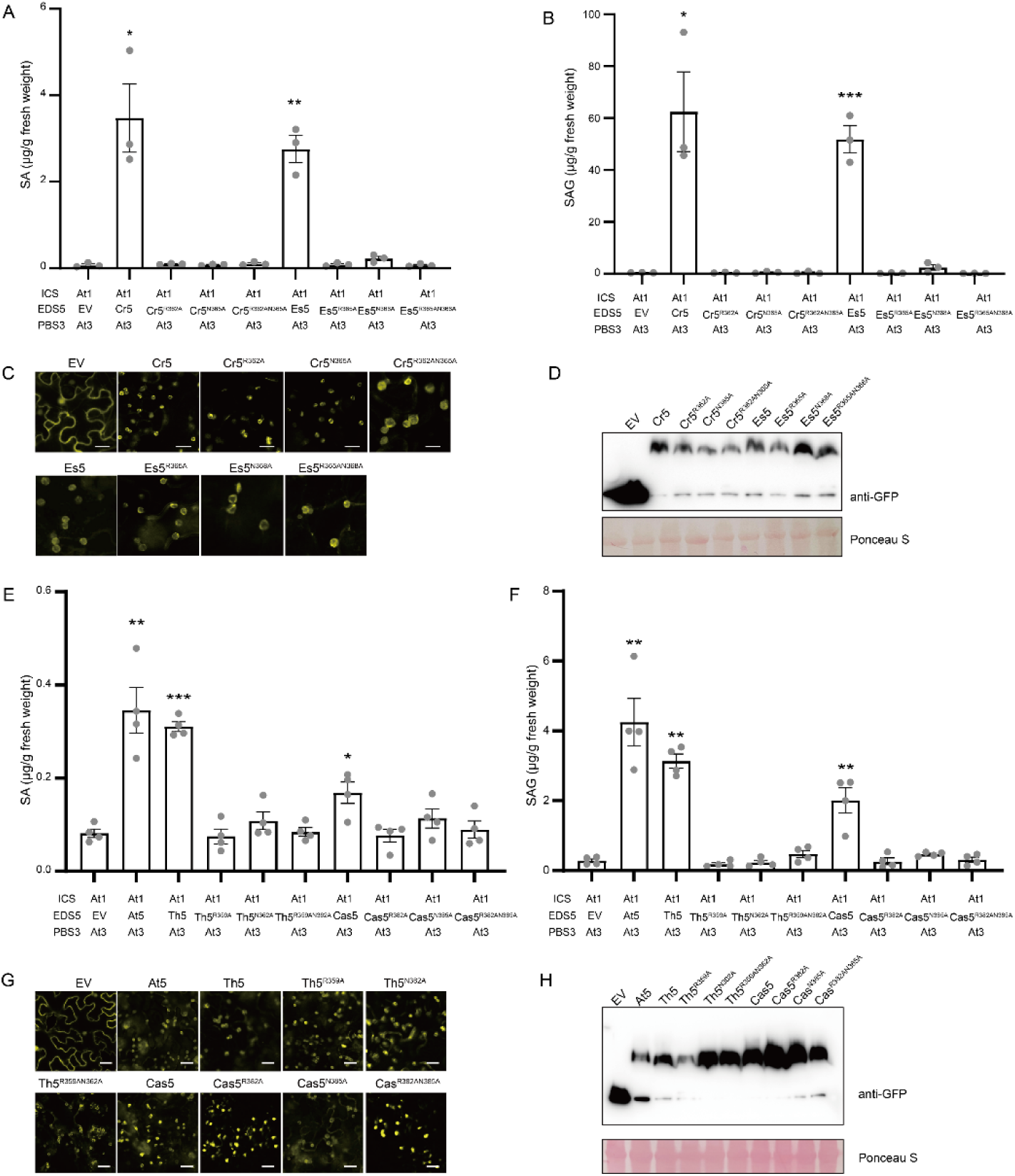
Conserved amino acids in the predicted IC-binding pocket of EDS5 are essential for SA biosynthesis. *Agrobacterium* strains carrying *EDS5-YFP* (or mutated *EDS5-YFP*) from *A. thaliana*, *C. rubella*, *E. salsugineum*, *T. hassleriana*, and *C. spinosa* were co-infiltrated with *Agrobacterium* strains carrying *A. thaliana ICS1-Myc* and *PBS3-YFP* into leaves of 4 to 6-week-old *N. benthamiana* plants. (**A**, **B**, **E**, and **F**) SA and SAG levels in leaf extracts were measured at 5 dpi. EV represents the empty vector control. Data represent means ± SEM from 4 biological replicates. Asterisks indicate significant differences compared to EV (\**P <* 0.05, \*\**P <* 0.01, and \*\*\**P* < 0.001, two-tailed Student’s t-tests). (**C** and **G**) *Agrobacterium* strains carrying *EDS5-YFP* were infiltrated into *N. benthamiana* leaves. EDS5 subcellular localization in epidermal cells expressing EDS5-YFP was analyzed by confocal microscopy at 2 dpi. Scale bars, 20 µm. (**D** and **H**) *Agrobacterium* strains carrying EDS5-YFP were infiltrated into *N. benthamiana* leaves. Total protein was extracted from the infiltrated leaves at 2 dpi, and EDS5 protein expression was assessed by immunoblotting using an anti-GFP antibody.

**Figure S16.**
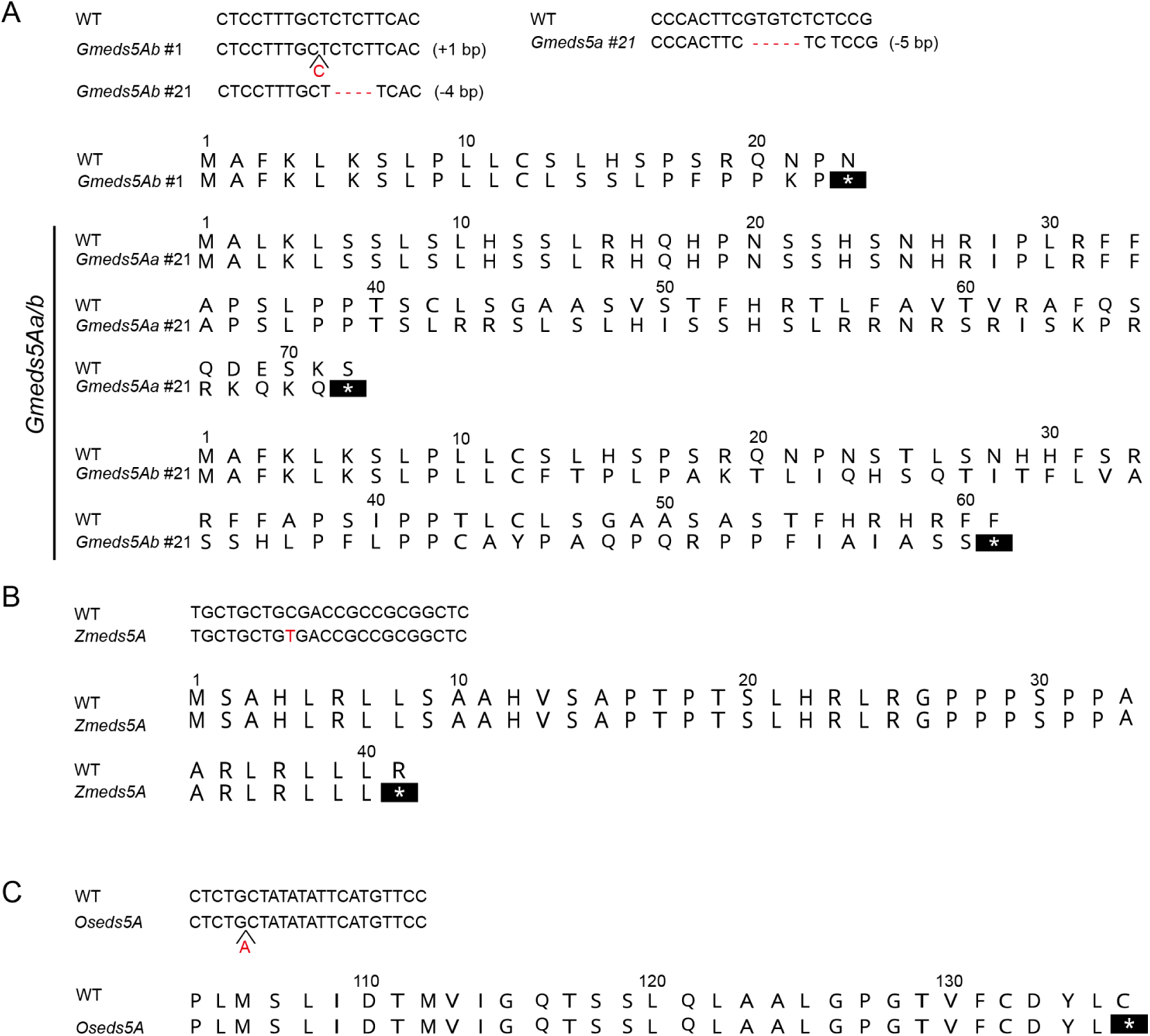
Characterization of *eds5A* mutants from soybean, maize, and rice. (**A**) Mutation sites and homozygosity in *Gmeds5A* mutants were verified by Sanger sequencing. Insertions and deletions led to frameshifts and the introduction of premature stop codons. (**B**) Mutation sites and homozygosity in *Zmeds5A* mutants were verified by Sanger sequencing. The mutation resulted in a premature stop codon. (**C**) Mutation sites and homozygosity in *Oseds5A* mutants were verified by Sanger sequencing. The insertion caused a frameshift, leading to a premature stop codon.

**Figure S17.**
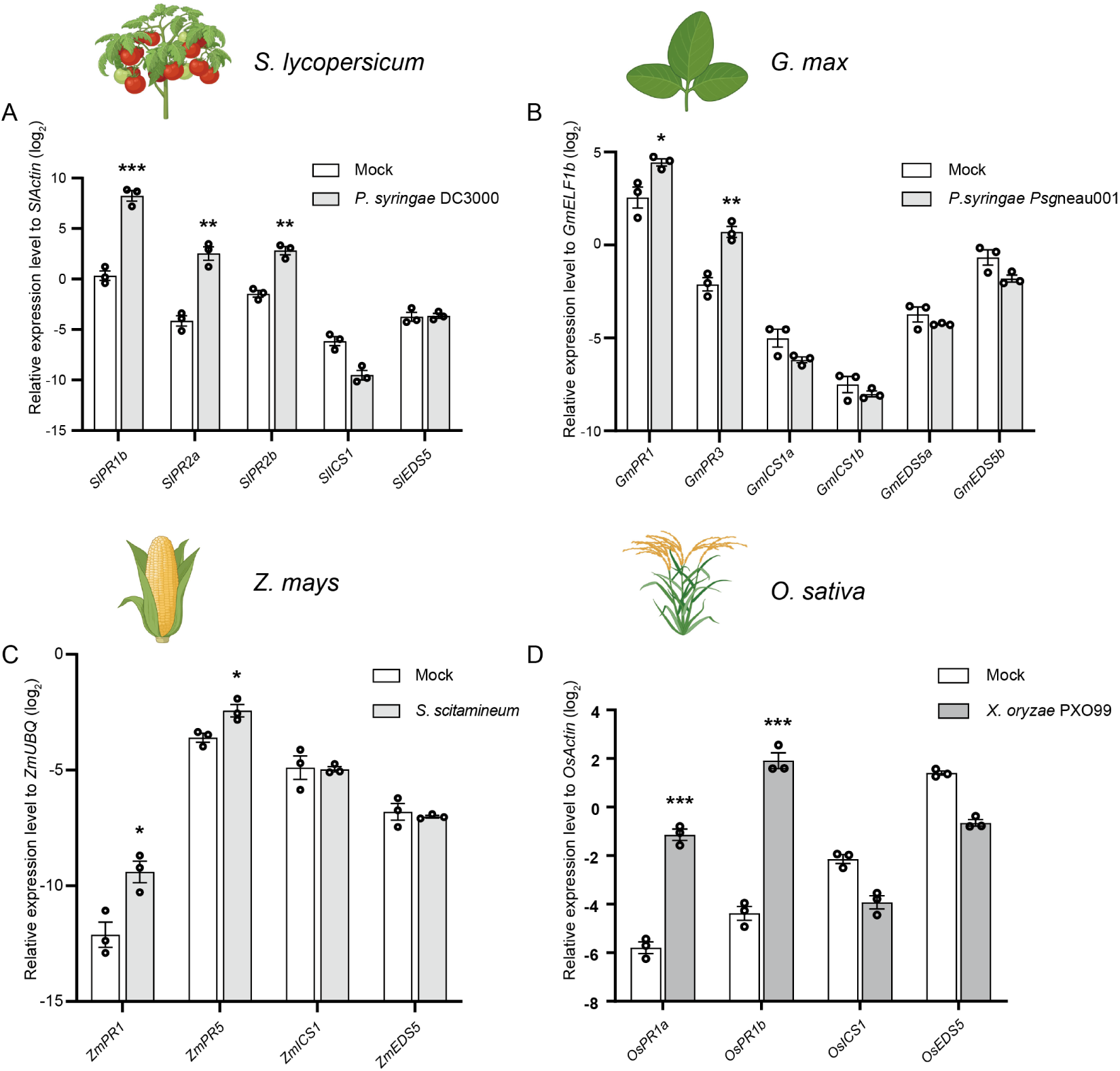
*ICS* and *EDS5A* expression is not induced following pathogen infection in *tomato*, soybean, maize, and rice. (**A**) Expression levels of *SlPR1b*, *SlPR2a*, *SlPR2b*, *SlICS1*, and *SlEDS5A* in 3-week-old tomato (Moneymaker) leaves at 24 h post-infiltration with *P. syringae* DC3000 (OD_600_ = 0.01) or mock, as determined by qRT-PCR. Values represent relative log_2_ expression levels normalized to *SlActin*. (**B**) Expression levels of *GmPR1*, *GmPR3*, *GmICS1a*, *GmICS1b*, *GmEDS5Aa*, and *GmEDS5Ab* in 2-week-old soybean (Williams82) leaves at 48 h post-infiltration with *P. syringae Psg*neau001 (OD_600_ = 0.5) or mock, as determined by qRT-PCR. Values represent relative log_2_ expression levels normalized to *GmELF1b*. (**C**) Expression levels of *ZmPR1*, *ZmPR5*, *ZmICS1*, and *ZmEDS5A* in 1-week-old maize (B73) leaves at 48 h post-injection with *Sporisorium scitamineum* (OD_600_ = 1.0) or mock, as determined by qRT-PCR. Values represent relative log_2_ expression levels normalized to *ZmUBQ*. (**D**) Expression levels of *OsPR1a*, *OsPR1b*, *OsICS1*, and *OsEDS5A* in 2-week-old rice (Kitaake) leaves at 48 h post-infiltration with *Xathomonas oryza* PXO99 (OD_600_ = 0.01) or mock, as determined by qRT-PCR. Values represent relative log_2_ expression levels normalized to *OsActin*. (**A-D**) Data represent means ± SEM from 3 biological replicates. Asterisks indicate significant differences compared to mock (*P < 0.05, **P < 0.01, and ***P < 0.001, two-tailed Student’s t-tests).

**Figure S18.**
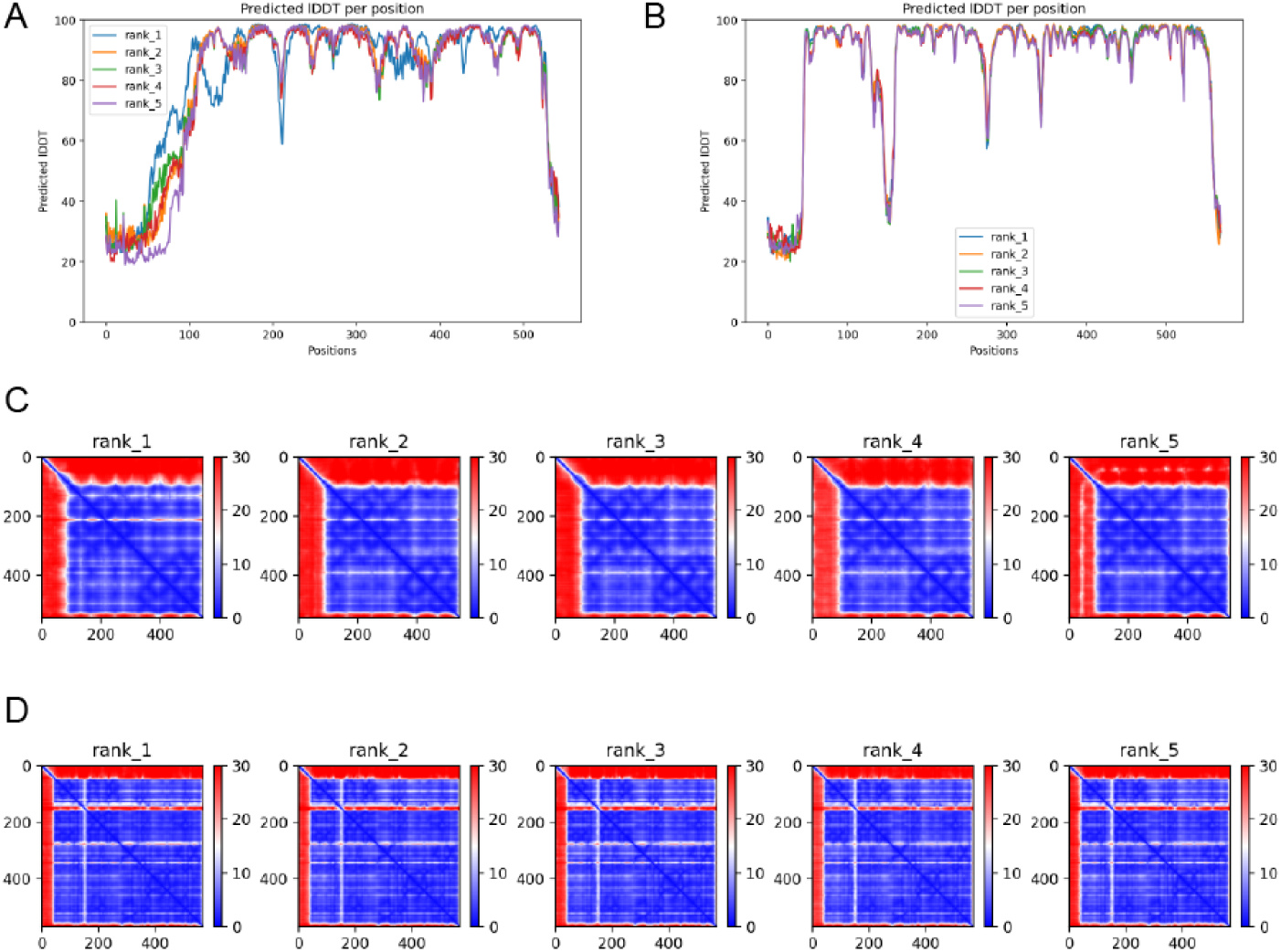
Structure prediction statistics. (A–B) pLDDT scores mapped to each amino acid residue in AtEDS5 (A) and AtICS1 (B). (C–D) PAE scores for AtEDS5 (C) and AtICS1 (D).

